# Antigen presentation by tumor-associated macrophages drives tumor-infiltrating T cells from a progenitor exhaustion state to terminal exhaustion

**DOI:** 10.1101/2023.08.11.553007

**Authors:** Jessica Waibl Polania, Alexandra Hoyt-Miggelbrink, William H. Tomaszewski, Lucas P. Wachsmuth, Selena J. Lorrey, Daniel S. Wilkinson, Emily Lerner, Karolina Woroniecka, John B. Finlay, Katayoun Ayasoufi, Peter E. Fecci

## Abstract

Whereas terminally exhausted T (Tex_term) cells retain anti-tumor cytotoxic functions, the frequencies of stem-like progenitor exhausted T (Tex_prog) cells better reflect immunotherapeutic responsivity. Here, we examined the intratumoral cellular interactions that govern the transition to terminal T cell exhaustion. We defined a metric reflecting the intratumoral progenitor exhaustion-to-terminal exhaustion ratio (PETER), which decreased with tumor progression in solid cancers. Single cell analyses of Tex_prog cells and Tex_term cells in glioblastoma (GBM), a setting of severe T cell exhaustion, revealed disproportionate loss of Tex_prog cells over time. Exhaustion concentrated within tumor-specific T cell subsets, with cognate antigen exposure requisite for acquisition of the Tex_term phenotype. Tumor-associated macrophages (TAM) - not tumor cells - were the primary source of antigenic exposure governing the Tex_prog to Tex_term transition. TAM depletion increased frequencies of Tex_prog cells in multiple tumor models, increased PETER, and promoted responsiveness to αPD-1 immunotherapy. Thus, targeting TAM – T cell interactions may further license checkpoint blockade responses.

## INTRODUCTION

Brain tumors, both primary and metastatic, continue to herald poor survival^1,2^. Immunotherapies, such as immune checkpoint blockade (ICB) and chimeric antigen receptor (CAR) T cells, hold promise for improving outcomes, but remain only marginally effective within the intracranial confines^3,4^. This has been especially true in the case of primary malignant brain tumors, such as glioblastoma (GBM)^5^. An established contributor to immunotherapeutic failures in GBM and other solid cancers is tumor-imposed T cell dysfunction, particularly exhaustion^6–13^. T cell exhaustion is a differentiation state characterized by progressive and hierarchical loss of effector functions following chronic antigen exposure^14–17^. T cell exhaustion is severe in GBM^6^, making GBM an important model tumor for studying the dynamics of and contributors to T cell exhaustion.

Exhausted T cells are divided into two subsets: progenitor exhausted T (Tex_prog) cells and terminally exhausted T (Tex_term) cells, the latter of which are also frequently referred to as “terminally differentiated” or “terminal subsets” since Tex_term cells typically still maintain cytotoxic function. We will use the exhaustion nomenclature (Tex_prog and Tex_term) throughout this manuscript^18–22^. Tex_prog describes a stem-like state, characterized by the expression of PD-1 and TCF1 (surrogate Ly108), which retains T cell polyfunctionality and is immunotherapy-responsive^21^. Tex_prog cells may subsequently differentiate toward Tex_term cells, which are characterized by high expression of TOX, PD-1 and TIM3. Tex_term cells have greater cytotoxic capability, but are short-lived and non-replenishing^15,16,21^. Promoting the maintenance of Tex_prog is an important consideration for generating an immunotherapy-responsive tumor microenvironment (TME)^18,21,23,24^. Ultimately, then, while Tex_term cells retain important anti-tumor cytotoxic functions, it is the relative renewal of the stem-like progenitor exhaustion state (Tex_prog) that better reflects immunotherapeutic responsivity. Despite this, the cellular interactions in a tumor microenvironment (TME) governing the transition between Tex_prog and Tex_term are not yet entirely established.

Tumor cells are frequently regarded as direct sources of T cell dysfunction within the TME, and the presumed source of the chronic antigen exposure promoting T cell exhaustion^25,26^. However, antigen presentation by infiltrating myeloid cells, such as tumor associated macrophages (TAM) and dendritic cells (DC) may also contribute to driving T cell exhaustion^27,28^.

Here, we characterized the progression of T cells from Tex_prog to Tex_term state within the TME. We observed a disproportionate loss of Tex_prog cells over time in GBM, leading to a low progenitor exhaustion to terminal exhaustion ratio (PETER). Terminal exhaustion was concentrated within tumor-specific T cell subsets, with cognate antigenic exposure being requisite for acquisition of the Tex_term phenotype. TAM, not tumor cells, were the relevant source of antigenic exposure governing the Tex_prog to Tex_term transition; antigen cognate T cell interactions with MHC I^+^ TAM were required for the transition to Tex_term state. TAM depletion increased PETER across multiple tumor models, and depletion of MHC I on tumor-infiltrating TAM reduced the accumulation of Tex_term T cells, with the degree of Tex_term cell persistence correlating with residual TAM MHC I expression. Cell – cell interaction analsyes suggest secondary TAM – T cell receptor-ligand communications that may contribute to T cell progression to terminal exhaustion and to a resultant decline in PETER within the TME. TAM depletion enhanced αPD1 responses and tumor control. The highlighted role for TAM antigen presentation in driving the intratumoral Tex_prog to Tex_term transition advance our understanding of how exhaustion develops within the TME and suggest TAM – T cell interactions as a therapeutic target for further licensing checkpoint blockade responses

## RESULTS

### GBM-infiltrating CD8^+^ T cells demonstrate a decline in the Progenitor Exhaustion to Terminal Exhaustion Ratio (PETER) over time

To better examine the dynamics of T cell exhaustion in GBM and other solid tumor models, but to eliminate any variables that might be imposed by anatomic compartment, we implanted murine GBM, melanoma, and lung (CT2A, B16F0, and LLC, respectively) tumor cells intracranially into syngeneic C57BL/6 mice, harvesting tumors when mice became moribund. We began by evaluating the frequency of both progenitor (Tex_prog) (PD1^+^Ly108^+^TIM3^−^) and terminal (Tex_term) (PD1^+^Ly108^−^TIM3^+^) exhaustion in each model (**Figure 1A**). Consistent with our previous findings^6^, GBM tumors demonstrated a higher frequency of Tex_term cells and correspondingly lower frequency of Tex_prog cells than the brain metastasis models (**Figure 1B and Figure S1A**).

**Figure 1:**
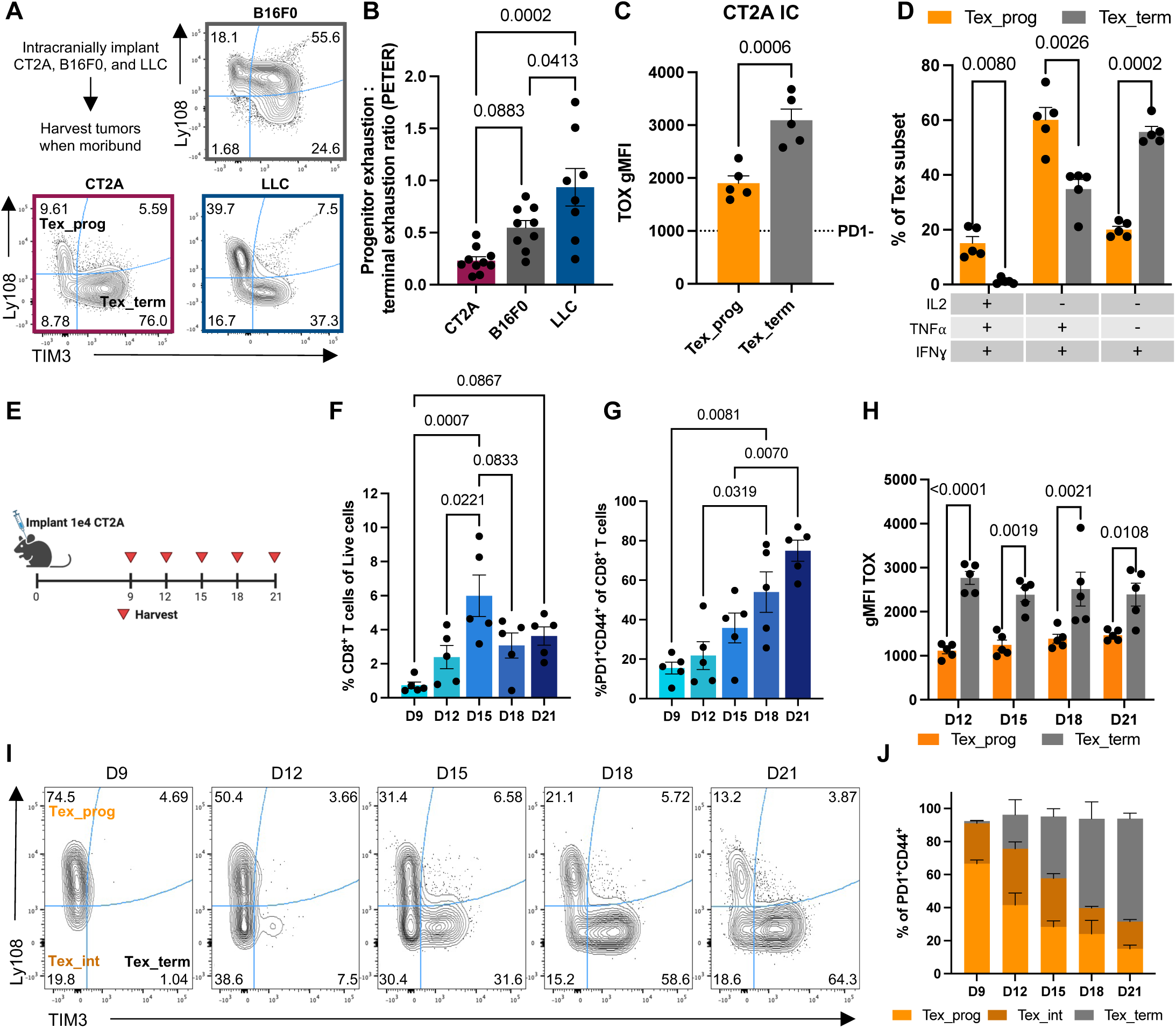
GBM-infiltrating CD8^+^ T cells demonstrate a decline in the Progenitor Exhaustion to Terminal Exhaustion Ratio (PETER) over time. (A) Representative contour plots of Tex_prog cells and Tex_term cells across tumor types. (B) PETER across CT2A, B16F0 and LLC intracranial tumors, harvested when mice were moribund. n = 8-10 mice/group. (C and D) Geometric mean fluorescence intensity (gMFI) of intracellular TOX expression (C) and quantification of TNFα/IFNγ/IL2 of Tex_prog cells and Tex_term cells in 10K CT2A at D18 (D). n = 5 mice. Representative of two independent experiments. (E) Experimental layout. 10K CT2A cells implanted D0, tumors were harvested every 3 days starting D9 until mice were moribund on D21. Created with Biorender. (F) number of CD8+ T cells per harvested tumor. n = 5 mice/timepoint. (G) Frequency of PD1^+^CD44^+^ of CD8^+^ T cells. n = 5 mice/timepoint. (H) Geometric mean fluorescence intensity (gMFI) of TOX of Tex_prog cells and Tex_term cells across timepoints with >100 total PD1^+^CD44^+^ CD8^+^ T cells. n = 5 mice/timepoint. (I and J) Representative plots showing Ly108 vs. TIM3 progression from D9 to D21, Tex_prog cells = top left, Tex_int cells = bottom left, Tex_term cells = bottom right (I) and quantification of Tex subsets of PD1^+^CD44^+^ CD8+ T cells over time (J). n = 5 mice/timepoint. All of the data are means ± SEMs; p-values are shown on individual plots as determined by 1-way ANOVA and post-hoc Tukey test (A, F, G), two-sided paired t-test (B) or RM 2-way ANOVA and post-hoc Tukey test (D and H). See also Figure S1.

Having found that GBM possessed a particularly low progenitor exhaustion to terminal exhaustion ratio (which we have coined “PETER”), we took this to indicate that GBM-infiltrating T cells may transition from Tex_prog to Tex_term at an especially high rate. We therefore selected GBM as an archetype tumor model to further examine the processes by which this transition occurs. To begin, we validated the expected transcriptional and functional differences across tumor-infiltrating Tex_prog cells and Tex_term cells from CT2A-bearing mice. We observed higher TOX expression in Tex_term cells compared to Tex_prog cells, with both subsets having higher expression than non-exhausted PD1^−^ CD8^+^ T cells, confirming the typical transcriptional landscape (**Figure 1C**). Likewise, upon PMA/Ionomycin stimulation *ex-vivo*, Tex_prog cells retained the ability to secrete IL2, IFNγ, and TNFα, whereas Tex_term cells predominantly secreted IFNγ alone (**Figure 1D**). Consistent with findings from other groups^22^, we observe only Tex_prog cells (and only at low levels) in the tumor-draining lymph node (tdLN), suggesting that the transition from Tex_prog to Tex_term is occurring later within the TME. Additionally, CD44^+^PD1^+^CD8^+^ T cells maintain a Ly108^+^TIM3^−^ Tex_prog phenotype and do not progress to Tex_term cells, even when they are trapped in the tdLN with FTY720 administration (**Figure S1C**).

We next examined the evolution of T cell exhaustion within the tumor over time. We harvested CT2A tumors every 3 days from day 9 (D9) through D21 post-implantation, where mice were moribund at D21 (**Figure 1E**). T cell infiltration was observed within the tumor at D9, with substantial increases seen by D15 (**Figure 1F**). Furthermore, CD8^+^ PD1^+^CD44^+^ T cells increased in frequency from <20% on day 9 to >75% by D21 (**Figure 1G**). Within the PD1^+^CD44^+^ subset, we identified Tex_prog cells and Tex_term cells as above, with transcriptional exhaustion confirmed via TOX expression **(Figure 1H**). Higher TOX levels were consistently seen in Tex_term cells than in Tex_prog cells at all timepoints in which both subsets were present (D12-21), confirming the anticipated pattern of expression (**Figure 1H**).

Tex_prog cells were readily identifiable by D9 and accounted for ∼60% of the PD1^+^CD8^+^ T cells found at the tumor, while Tex_term cells were notably absent at this early timepoint. The frequency of Tex_prog cells subsequently decreased over the course of tumor progression, accounting for less than 20% of PD1^+^CD8^+^ T cells by D21. (**Figures 1I and 1J**). Tex_term cells were identifiable starting D12 at low frequencies (∼10%). By D21, however, Tex_term cells accounted for ∼70% of PD1^+^CD44^+^ CD8^+^ T cells (**Figures 1I and 1J**), further revealing a drop in PETER over time.

### scRNAseq defines transcriptional heterogeneity amongst GBM-infiltrating Tex_prog cells and Tex_term cells

To better understand the requirements for the transition from Tex_prog to Tex_term, we first performed scRNAseq on sorted CD8^+^ T cells from glioma-bearing mice. Informed by our timecourse, we chose an early (D13) and a late (D19) timepoint for our analyses (**Figure 2A**). We identified six clusters of CD8^+^ T cells, including two separate terminal exhaustion populations, which we termed Tex_term_early and Tex_term_late. While 5 of the clusters were present at both timepoints (D13 and 19), Tex_term_early cells were present only at D13 (**Figures 2B and 2C**). Clusters were annotated based on their top differentially expressed genes and their expression of known T cell exhaustion genes, namely *Tcf7*, *Tox*, and *Pdcd1* **(Figure 2D and Figures S2A and 2B)**. Xcl1+_Tcm and Naïve-like clusters expressed memory/naïve T cell markers (*Sell, Il7r, Id3, Tcf7*), while Xcl1+_Tcm additionally expressed genes associated with activation and TCR signaling (*Xcl1, Klrk1, Nr4a2*) (**Figure 2D)**. The expression of *Pdcd1* and *Tox* on Tex_prog, Tex_int, Tex_term_early, and Tex_term clusters suggested that these were T cell exhaustion subsets (**Figure 2D and Figure S2B**). Tex_prog cells expressed effector (*Ifng, Tbx21*) and stem-like genes (*Cd28, Il7r, Tcf7, Tbx21*) and had a lower T cell exhaustion signature score (described by Im et al^18^) than Tex_int, Tex_term_early, and Tex_term_late cells (**Figures 2D and 2E**). This could suggest that the exhaustion program may not be fully established in Tex_prog cells. Next, Tex_int cells had increased expression of exhaustion-associated genes (*Pdcd1, Tox, Entpd1*) and a decrease in effector (*Klrg1*) and memory associated genes (*Tcf7, Il2*) (**Figure 2D**). Cytotoxicity-associated genes, such as *Gzmb, Gzma*, and *Gzmk* were expressed at the highest levels in Tex_term_early and Tex_term_late cells, consistent with previous characterization of terminal exhaustion^21^ (**Figure 2D**). Finally, Tex_term_late cells, predominantly found at the late timepoint and consistent with our time-course characterizations, expressed the highest levels of exhaustion-associated genes (*Pdcd1, Tox, Havcr2, Entpd1*) and lost *Tcf7* expression (**Figure 2D and Figure S2B**).

**Figure 2:**
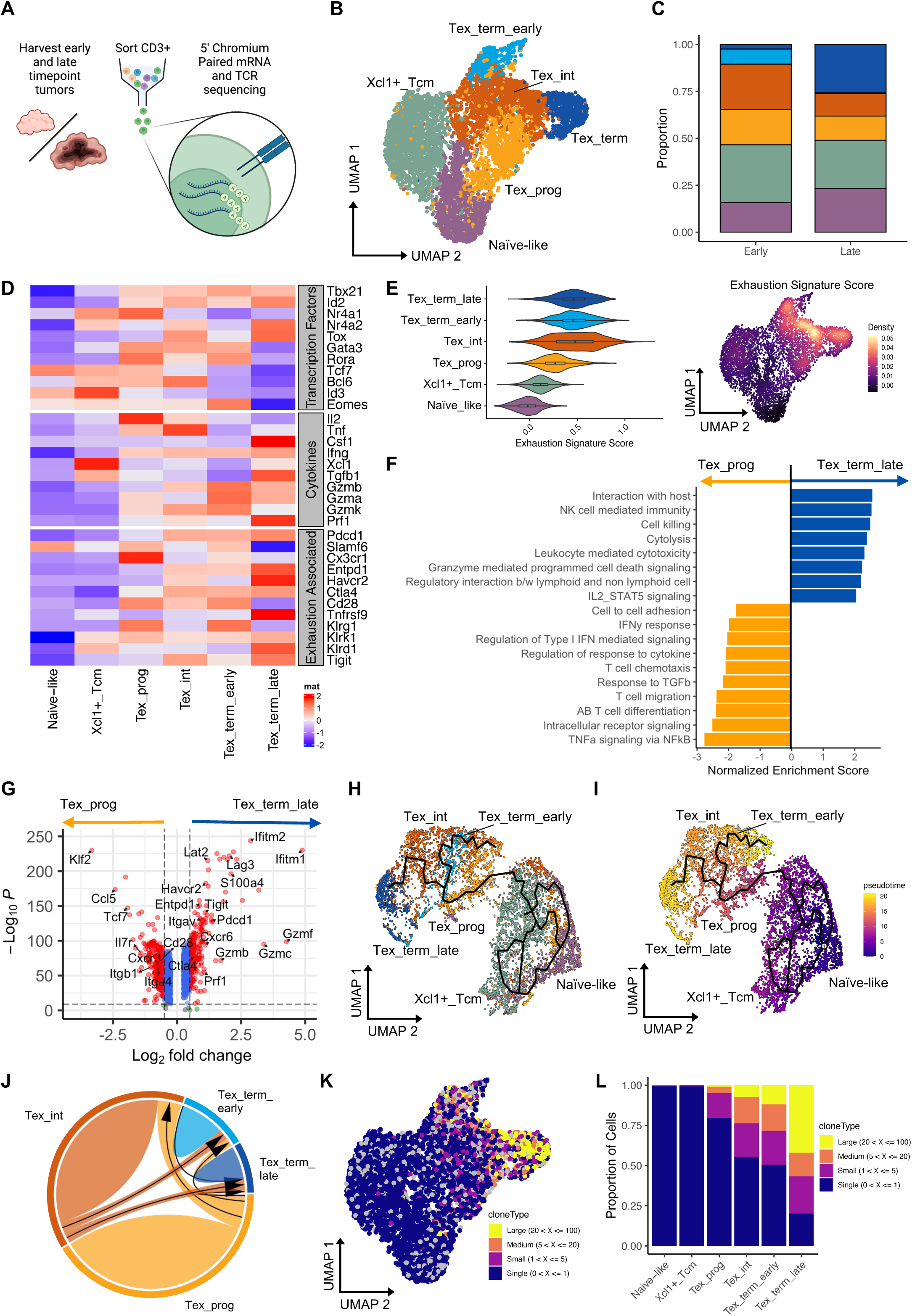
scRNAseq defines transcriptional heterogeneity amongst GBM-infiltrating Tex_prog cells and Tex_term cells. (A) Experimental layout for scRNAseq experiment. tumors were harvested at an early (D13) and late (D19) timepoint. Created with Biorender. (B) UMAP of 6 CD8 T cell clusters following QC. (C) Frequency of each cluster per timepoint. (D) Heatmap of exhaustion associated transcription factors, cytokines/secreted molecules, and surface markers. (E) AddModuleScore of exhaustion signature from Im et al., 2016, violin plot and density plot representations. (F) Significantly enriched pathways in Tex_prog and Tex_term from Hallmark, Reactome, Gene Ontology databases. (G) Relevant genes from enriched pathways in F. (H) Trajectory analysis overlaid onto clusters from B. (I) Trajectory analysis overlaid onto pseudotime. (J) Circle plot of overlapping clones across Tex_clusters. Tex_prog cells and Tex_int cells have overlapping clones with all other clusters. Tex_term_late cells and Tex_term_early cells do not have overlapping clones. (K) UMAP of cloneTypes across clusters. (L) Frequency of cloneTypes across clusters. See also Figure S2 and Table S1.

We subsequently employed pathway enrichment analysis for both Tex_prog and Tex_term_late cells, which represented the two extreme ends of the exhaustion pathway. This analysis revealed an abundance of pathways related to T cell migration, differentiation, cell-cell communication and response to stimuli (cytokine, chemokines) in Tex_prog cells (**Figures 2F and 2G**). This is in keeping with the current understanding of the Tex_prog state and its prevalence among T cells entering or recently entering tumor ^22,29^. In contrast, the Tex_term_late cluster was enriched for cytotoxic pathways, almost exclusively (**Figures 2F and 2G**). This highlights that Tex_prog cells, unlike Tex_term cells, likely remain in a signal-responsive state within the TME.

Lasty, we applied pseudotime analysis to our dataset to infer the temporal progression amongst the T cell exhaustion subsets. This analysis spatially separated the Naïve_like and Xcl1+_Tcm clusters from the other four clusters, supporting their identification as separate, non-exhausted lineages (**Figure 2H**). The pseudotime plot ultimately validated the progression from progenitor to terminal exhaustion through an intermediate population (Tex_int). However, two trajectories were determined, both beginning with Tex_prog (**Figure 2I**). While one ended with Tex_term_late, the other ended in Tex_term_early. This finding is likely simply highlighting differences in phenotypes for the predominant ending node at the early and late timepoints.

To further examine lineage relationships between the T cell exhaustion clusters, we utilized our paired T cell receptor sequencing (TCRseq) dataset. To this end, we observed TCR clonal overlap between predicted Tex clusters (**Figure 2J**). In combination with our pseudotime results, this analysis suggests a trajectory of shared TCR clones, beginning with Tex_prog cells, progressing through Tex_int cells and eventually ending with Tex_term cells. We next compared clonal expansion between clusters, where we observed the greatest frequency of “Large” expanded clones, (defined as between 20 and 100 cells with the same clone), in Tex_term_late cells, and the lowest such frequency in Tex_prog cells (**Figure 2K and 2L**). Correspondingly, Tex_prog cells had the greatest diversity index score, as measured by the Shannon and Inverse Simpson indices, which has often been associated with response to immunotherapy (**Figure S2C**)^30^. Moreover, when we plotted the 10 most expanded clones on a UMAP and divided the analysis by timepoint, we observed that the expanded clones overwhelmingly resided in Tex_int, Tex_term_early and Tex_term_late cells, but not Tex_prog cells (**Figure S2D**). This suggests that clonal expansion in response to TCR stimulation closely reflects T cell exhaustion progression.

### Tumor cells are not a necessary source of the TCR stimulation mediating Tex_prog to Tex_term transition

We hypothesized that antigen-specific TCR activation would be required for progression of Tex_prog to Tex_term within the TME, and thus that Tex_term would be concentrated amongst tumor-specific T cells. To test this directly, we employed a model in which CT2A gliomas were engineered to express the antigen TRP2 (See STAR Methods). We then adoptively transferred either tumor-specific TRP2 TCR-transgenic CD8^+^ T cells (TRP2-TCR) or irrelevant OVA antigen-specific (OT1) CD8^+^ T cells into CT2A-TRP2-bearing mice (**Figure 3A**). All T cells were activated *ex vivo* prior to transfer. Two weeks later, tumors were harvested, and adoptively transferred T cells were identified as TRP2 tetramer^+^CD45.1^+^ (TRP2-TCR) or OVA tetramer^+^ (OT1) and evaluated for exhaustion phenotypes. Recovery of TRP2-TCR T cells from tumor was better than recovery of OT1 T cells (**Figure 3B**). Recovered TRP2-TCR T cells also demonstrated higher acquisition of a Tex_term phenotype than OT1 T cells, which only minimally acquired PD1 expression (**Figures 3C and 3D and Figure S3A**). Unlike OT1 T cells, transferred TRP2-TCR T cells developed into predominantly Tex_int cells and Tex_term cells, similar to endogenous tumor-specific T cells (TRP2 tetramer^+^CD45.2^+^) (**Figure S3B)**. Taken together, these data suggest that T cells in the TME must encounter their cognate antigen to progress along the exhaustion pathway and complete the transition to Tex_term.

**Figure 3:**
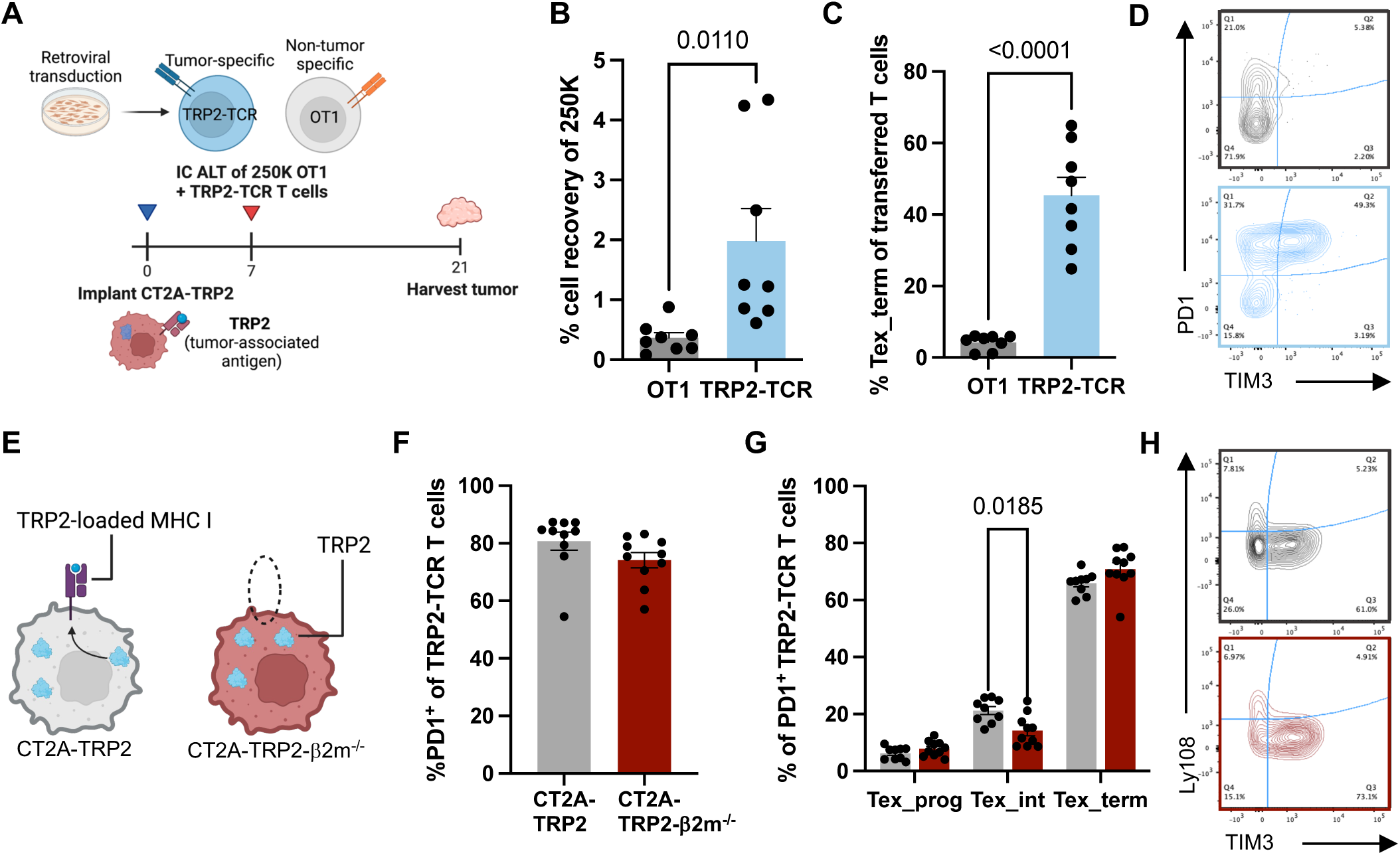
Tumor cells are not a necessary source of the TCR stimulation mediating Tex_prog to Tex_term transition. (A) Experimental layout. 2.5 × 10^5^ *ex-vivo* activated and transduced OT1 and TRP2-TCR T cells were intracranially implanted into D7 CT2A-TRP2-bearing mice. Tumors were harvested 14 days later. (B) Frequency of implanted cells recovered on D21. n = 7 mice/group. (C and D) Frequency of PD1^+^ CD8^+^ T cells in OT1 vs. TRP2-TCR (C) and representative contour plot PD1 and TIM3 expression of OT1 and TRP2-TCR T cells 14 days post-transfer (D). n = 7 mice/group. (E) Experimental layout. 5.0 × 10^4^ CT2A-TRP2 or CT2A-TRP2-β2m^−/-^ cells were implanted intracranially and harvested when mice were moribund. Created with Biorender. (F) Frequency of PD1^+^ TRP2-TCR T cells between groups. n = 10 mice/group. (G and H) Frequency of Tex subset of PD1^+^TRP2-Tetramer^+^ CD8^+^ T cells and representative contour plot (G) and representative contour plot of Ly108 and TIM3 expression between groups (H). n = 10 mice/group. All of the data are means ± SEMs; p-values are shown on individual plots as measured by two tailed t-test (B, C, F) or RM 2-way ANOVA (G). See also Figure S3.

As tumor antigen-specific TCR stimulation appeared to be necessary for exhaustion progression, we first supposed that the relevant antigen-cognate interaction would be between T cells and tumor cells. To evaluate this, we compared the progression of T cell exhaustion *in vivo* within CT2A-TRP2 and CT2A-TRP2-β2m^−/-^ tumors, where the latter constitutes a cell line where β2m is genetically deleted using CRISPR-Cas9 (See STAR Methods). Genetic deletion of β2m leaves the tumor without the capacity to build functional MHC I molecules, and thus without the capacity to present antigen to T cells (**Figure 3E and Figure S3C**). We observed no difference in the frequency of PD1^+^ TRP2-TCR T cells or in the frequency of T cell exhaustion subsets (Tex_prog or Tex_term) in the absence of tumor MHC I expression and antigen presentation (**Figures 3F, 3G, and 3H**). Of note, the extent of T cell infiltration was comparable between MHC I expressing and non-expressing tumors (**Figure S3D**).

### TAM are abundant cross-presenting antigen-presenting cells (APC) within the intracranial TME

With tumor cells seemingly eliminated as the source of exhaustion-relevant antigen-cognate interactions with T cells, we next sought to narrow down additional candidate antigen presenting populations. We therefore evaluated other sources of potentially determinant antigen-cognate interactions within the TME. Such relevant APC might include monocytes, TAM, microglia, and/or dendritic cells (DC)^31^. Using a scRNAseq dataset of CD45^+^ cells sorted from CT2A tumors^32^, we identified predicted sources of MHC I cell-cell interactions in the TME. We isolated a component of the dataset that included the above APC types and CD8^+^ T cells, then conducted cell-cell interaction analysis using the CellChat pipeline^32,33^ (Schematic in **Figure 4A**). MHC I signaling was a highly predicted interaction between T cells and all potential APC based on differentially expressed receptor/ligand pairs (**Figure 4B**). This analysis revealed the greatest *number* of predicted interactions between CD8^+^ T cells and Apoe^+^ TAM, while the *strength* of predicted interactions was similar across APC (**Figure 4C and Figure S4A**). TAM greatly outnumber other APC in the CT2A TME, comprising ∼40% of tumor-infiltrating immune cells (**Figure 4D**). Combined, these results suggested that the relevant antigen-cognate interactions promoting exhaustion progression within the TME are occurring between CD8^+^ T cells and APC (and not tumor), with TAM being the the most abundant source of antigen.

**Figure 4:**
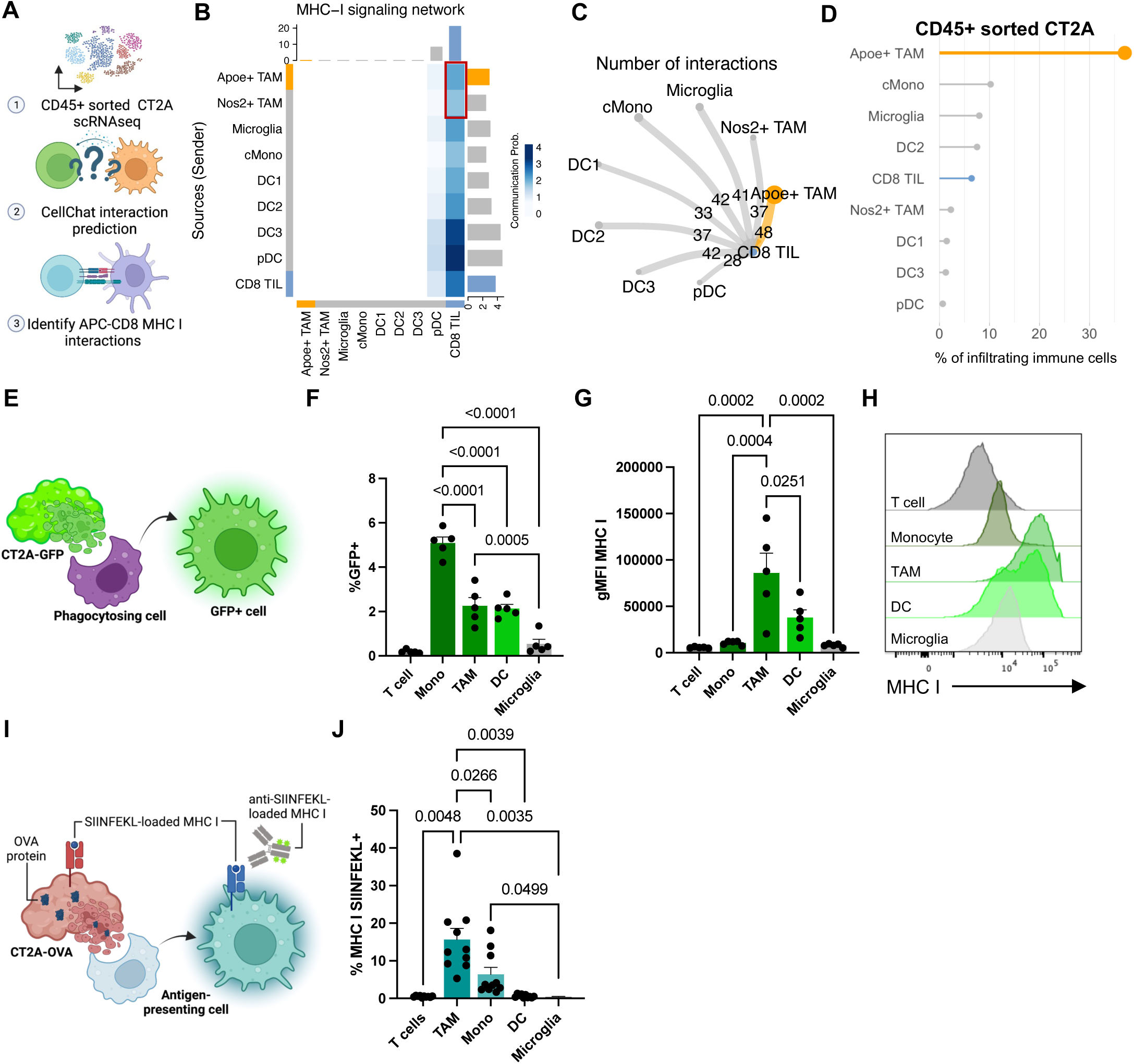
TAM are abundant cross-presenting antigen-presenting cells (APC) within the intracranial TME. (A) Experimental layout. CD45^+^ sorted scRNAseq from Tomaszewski et al., 2022 was analyzed with CellChat to determine which APC had predicted MHC I interactions with CD8 T cells. Created with Biorender. (B) Heatmap showing communication probability of MHC I interactions in the TME. All APC were predicted to interact with CD8 TIL through MHC I. (C) Circle plot displaying number of interactions (thickness of arrow/number) across APC populations. (D) Frequency of infiltrating immune cells from Tomaszewski et al., 2022 dataset. (E) Experimental layout. 2.5 × 10^4^ CT2A-GFP was implanted intracranially and harvested when mice were moribund. GFP^+^ APC were evaluated. Created with Biorender. (F) Frequency of GFP^+^ T cells (CD3^+^), TAM (CD45^hi^CD11b^+^F4/80^+^), DC (CD11b^+^MHC II^+^F4/80^−^) and Microglia (CD45^hi^CD11b^+^F4/80^+^). n = 5 mice. (G and H) gMFI of MHC I of APC populations from F (G) and representative histogram (H). n = 5 mice. (I) Experimental layout. 1.0 × 10^5^ CT2A-OVA cells were implanted intracranially and harvested when mice were moribund. (J) Frequency of SIINFEKL-loaded MHC I of T cells, TAM, Monocytes (CD11b+F480^int^Ly6C^hi^), DC, and Microglia. n = 9 mice. p-values are plotted on individual graphs. Statistical significance was determined by 1-way ANOVA (F, G, J) with post-hoc Tukey’s test for multiple comparisons. All data are plotted as mean ± SEM. See also Figure S4.

To investigate the relevant antigen-presenting functions of the various APC subtypes within CT2A, we directly evaluated their relative *in vivo* phagocytic activity within the TME. To this end, we implanted GFP-expressing CT2A tumors and evaluated the frequency of GFP positivity within intratumoral APC when mice were moribund (schematic in **Figure 4E**). We focused on monocytes (CD11b^+^F4/80^+^Ly6C^hi^), TAM (CD11b^+^F4/80^+^MHC II^+^), DC (F4/80^−^CD11c^+^MHC II^+^), and microglia (CD45^int^CD11b^+^F4/80^+^) (**Figure S4B**). We found that monocytes, TAM, and DC all took up GFP (**Figure 4F**). After phagocytosing exogenous antigen, APC must load and cross-present the antigen via MHC I to interact with CD8^+^ T cells^34^. Therefore, we next probed for differential MHC I expression as a surrogate for subsequent tumor antigen cross-presentation. TAM and DC both had higher MHC I on their surface than other APC populations (**Figures 4G and 4H**).

To more directly evaluate the cross-presentation capacity of intratumoral APC, we implanted OVA-expressing CT2A tumors and subsequently assessed the various APC populations by flow cytometry for the presence of SIINFEKL-loaded MHC I (Schematic in **Figure 4I**). Among APC, we observed the greatest frequency of SIINFEKL-loaded MHC I on TAM and their monocyte precursors (**Figure 4J**). Combining our data, TAM appeared to exhibit the greatest potency for both phagocytosing and cross-presenting tumor antigen within the TME.

### TAM mediate Tex_prog to Tex_term transition in GBM and other solid tumor models

To better establish the requirement for TAM in the progression from Tex_prog to Tex_term, we implanted CT2A tumors into CCR2^−/-^ mice. CCR2^−/-^ mice exhibit impaired monocyte trafficking toward sites of inflammation^35^, therefore establishing a TME that is relatively deficient in TAM, which are monocyte-derived. To further the extent of TAM depletion in this model, however, we also administered tumor-bearing CCR2^−/-^ mice an anti-CSF1R antibody, which eliminates differentiated TAM (schematic in **Figure 5A**)^36^. We observed efficient depletion of TAM with this approach (**Figure S5A**). There was an increase in PETER observed in the CCR2^−/-^ setting (**Figures 5B and 5C**), suggesting a relative failure to progress from Tex_prog to Tex_term. We observed a strong positive correlation between the frequency of Tex_term cells and the frequency of TAM remaining within the TME, as Tex_term cell proportions dropped substantially with improved TAM depletion (**Figure 5D**). We did not observe similar alterations to PETER when we depleted DC (**Figure 5E and 5F**) or microglia (**Figure 5G and 5H) (Figures S5B-5E**), further suggesting TAM as the pertinent APC population for driving the progression of T cell exhaustion in this model.

**Figure 5:**
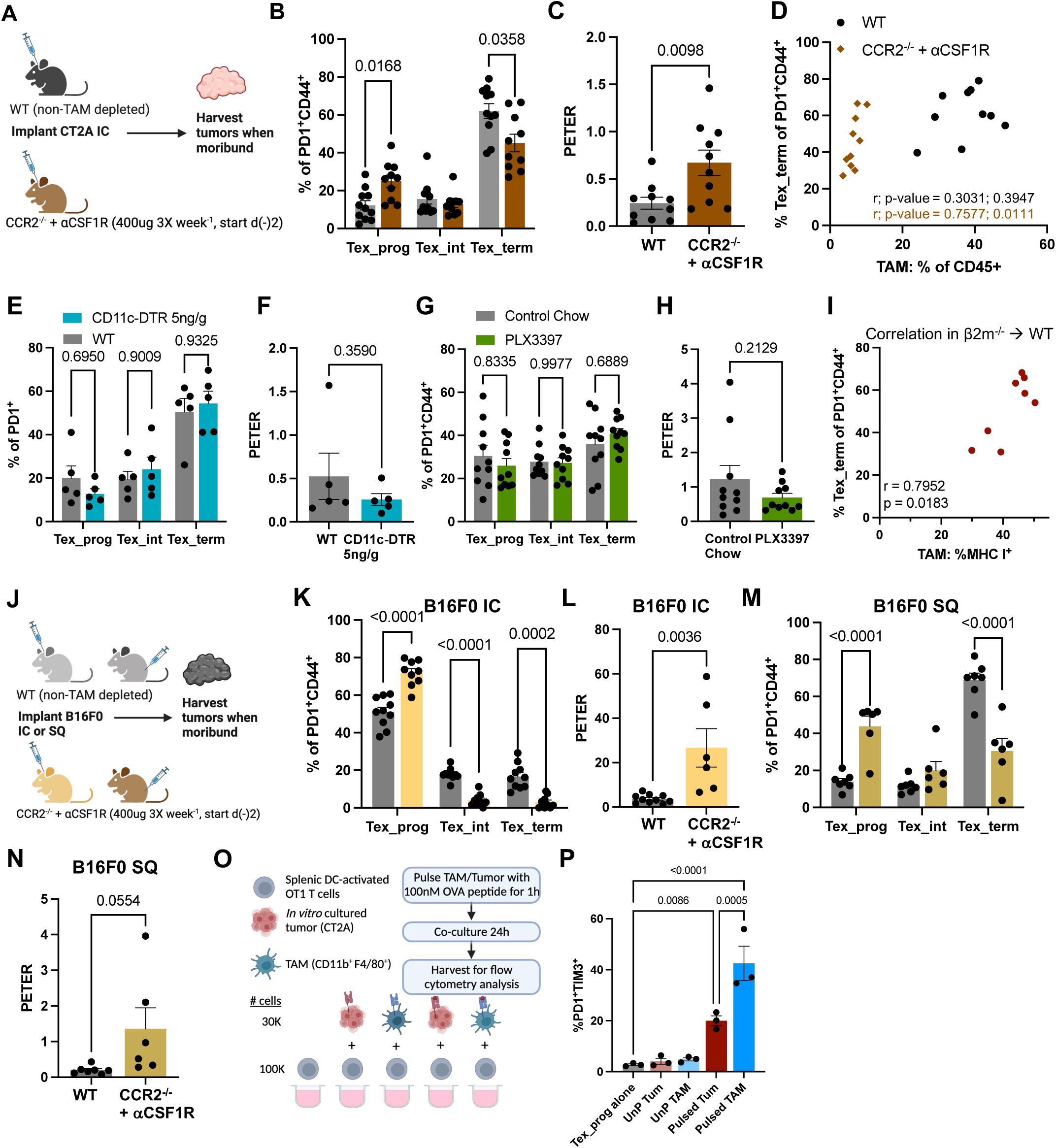
TAM mediate Tex_prog to Tex_term transition in GBM and other intracranial solid tumor models. (A) Experimental layout. TAM and monocytes were depleted in CCR2^−/-^ mice with αCSF1R depletion antibody. 1.0 × 10^4^ CT2A cells were implanted intracranially and harvested when mice were moribund. Created with Biorender. (B and C) Frequency of Tex_subsets of PD1^+^CD44^+^ (B) and PETER between WT and TAM depleted groups (C). Data pooled from 2 independent experiments. n = 10-11 mice/group. (D) Simple linear correlation of %Tex_term cells of PD1^+^CD44^+^ CD8^+^ T cells vs. % TAM of CD45^+^ cells. Data pooled from 2 independent experiments. n = 11 mice/group. (E and F) CD11c^+^ DC were depleted with 5ng/g DTR injections on D8, D11, D14, D17 post CT2A-implantation. Frequency of T cell exhaustion subsets (E) and and PETER between WT and DC depleted groups (F). n = 5 mice/group. (G and H) Microglia were depleted with 290mg/kg PLX3397 free-fed chow starting 1 week prior to CT2A implantation. Frequency of T cell exhaustion subsets (G) and PETER between WT and microglia depleted groups (H). n = 10 mice/group. (I) Pearson correlation of frequency of Tex_term cells vs. Frequency of MHC I+ TAM in β2m^−/-^ chimera mice implanted with CT2A. (J) Experimental layout. TAM and monocytes were depleted in CCR2^−/-^ mice with αCSF1R depletion antibody. 1.0 × 10^3^ B16F0 cells were implanted intracranially or 1.0 × 10^5^ B16F0 cells were implanted subcutaneously and harvested when mice were moribund. Created with Biorender. (K and L) Frequency of Tex_subsets of PD1^+^CD44^+^ B16F0 IC (K) and PETER between WT and TAM depleted groups (L). n = 9-10 mice/group. (M and N) Frequency of Tex subsets of PD1^+^CD44^+^ B16F0 SQ (M) and PETER between WT and TAM depleted groups (N). Data pooled from 2 independent experiments. n = 6-7 mice/group. (O) Experimental layout. Activated OT1 T cells were co-cultured with un-pulsed or peptide pulsed TAM or tumor cells for 24h. (P) Frequency of PD1^+^TIM3^+^ OT1 T cells when co-cultured with TAM or tumor. n = 3 biological replicates. Data are representative of 2 independent experiments. p-values are plotted on individual graphs. Statistical significance was determined by two-sided unpaired t test (C, F, H, I, L, N), 1-way ANOVA with post-hoc Tukey’s test (P) or RM-2-way ANOVA with post-hoc Tukey’s test for multiple comparisons (B, E, G, K, M). All data are plotted as mean ± SEM. See also Figure S5.

With these observations, we next evaluated the role of MHC I presentation by TAM in driving progression of T cell exhaustion. To this end, we generated β2m bone marrow chimeras (β2m^−/-^ donors into WT recipients), wherein hematopoietic cells (including hemaptopoietically-derived APC) lacked β2m and antigen presenting capacity (see STAR Methods). While we continued to observe low levels of MHC I expression on TAM in this model, (likely as a result of TAM trogocytosis of MHC I from surrounding β2m-replete cells^37^) (**Figures S5F and 5G**), we observed a positive correlation between persisting MHC I^+^ on TAM and the frequency of Tex_term (**Figure 5I**).

These findings in CT2A raised the question as to whether the role of TAM in driving exhaustion progression was exclusive to GBM and/or to the intracranial environment. To assess this, we employed the same TAM depletion approach in both orthotopic and intracranial murine models of melanoma (schematic in **Figure 5J**). As before, we saw efficient depletion of TAM (**Figures S5H and 5I**). In both peripheral and intracranial melanoma models, we observed an increase in PETER within the TME in the setting of TAM depletion (**Figures 5K-5N**).

To better isolate the impact of TAM antigen presentation we employed an *in vitro* co-culture system, where we co-cultured activated OT1 T cells, TAM, and tumor cells. OT1 T cells were activated for 48h with OVA peptide-pulsed splenic DC prior to co-culture (as described by Wu *et al*.^38^) to initiate a Tex_prog phenotype. TAM were sorted from IC CT2A tumors. Both TAM and cultured tumor cells were pulsed with OVA peptide for 1hr before co-culture with Tex_prog OT1 T cells. We facilitated MHC I-TCR interactions, specifically, by pulsing with an MHC I-restricted peptide (SIINFEKL). Neither TAM nor tumor cells possessed endogenous expression of OVA (**Figure 5O**). OT1 T cells had increased expression of PD1 and TIM3 only in the conditions where cognate MHC I-restricted antigen was presented by TAM or tumor cells. Co-culture with TAM led to higher transition to Tex_term than co-culture with tumor cells (**Figure 5P**). Additionally, we observed greater expression of the activation marker CD69 on OT1 T cells co-cultured with pulsed TAM compared to pulsed tumor (**Figure S5J**).

These findings implicate TAM as a primary source of tumor-antigen presentation mediating T cell exhaustion progression across multiple solid tumor types and locations. Taken together, these data suggest Tex_prog cells progress to Tex_term cells independently of tumor-derived antigen presentation, and instead rely heavily on antigen-cognate interactions between T cells and antigen cross-presenting APC, with TAM being the most prevalent of these in the TME.

### CellChat predicts secondary interactions between TAM and CD8^+^ T cells in mice and humans that may contribute to exhaustion progression

While TAM antigen cross presentation may serve as the initial interaction bringing TAM in contact with T cells within the TME, additional interactions are quite likely to play a role in driving exhaustion progression. To investigate this, we returned to the cell – cell interaction analysis from Figure 3 to further interrogate interactions between TAM and Tex_prog cells that might be contributors to Tex_prog progression to Tex_term (schematic in **Figure 6A**). Several TAM ligands or surface molecules were highlighted in our scRNAseq dataset, with pathways ranging from inhibitory (PD-L1, SPP1, GALECTIN) to stimulatory (CD86), to cell-cell adhesion (ICAM, JAM) to migration (CXCL), among others (**Figure 6B**). We further refined these possibilities by selecting receptors/ligands that were expressed in our scRNAseq dataset and specifically present in the Tex_prog cluster (**Figure 6C**). From these analyses, we observed that cell – cell adhesion (*Itgb1, Itgb2, Itga4*), migration (*Cxcr3*) and stimulatory molecules (*Cd28*) were particularly enriched in Tex_prog compared to Tex_term, consistent with the pathways identified in Figure 2F. Tex_term, by comparison, had greater expression of inhibitory receptors (*Pdcd1, Ctla4, Tigit*) and the chemokine receptor, *Cxcr6*.

**Figure 6:**
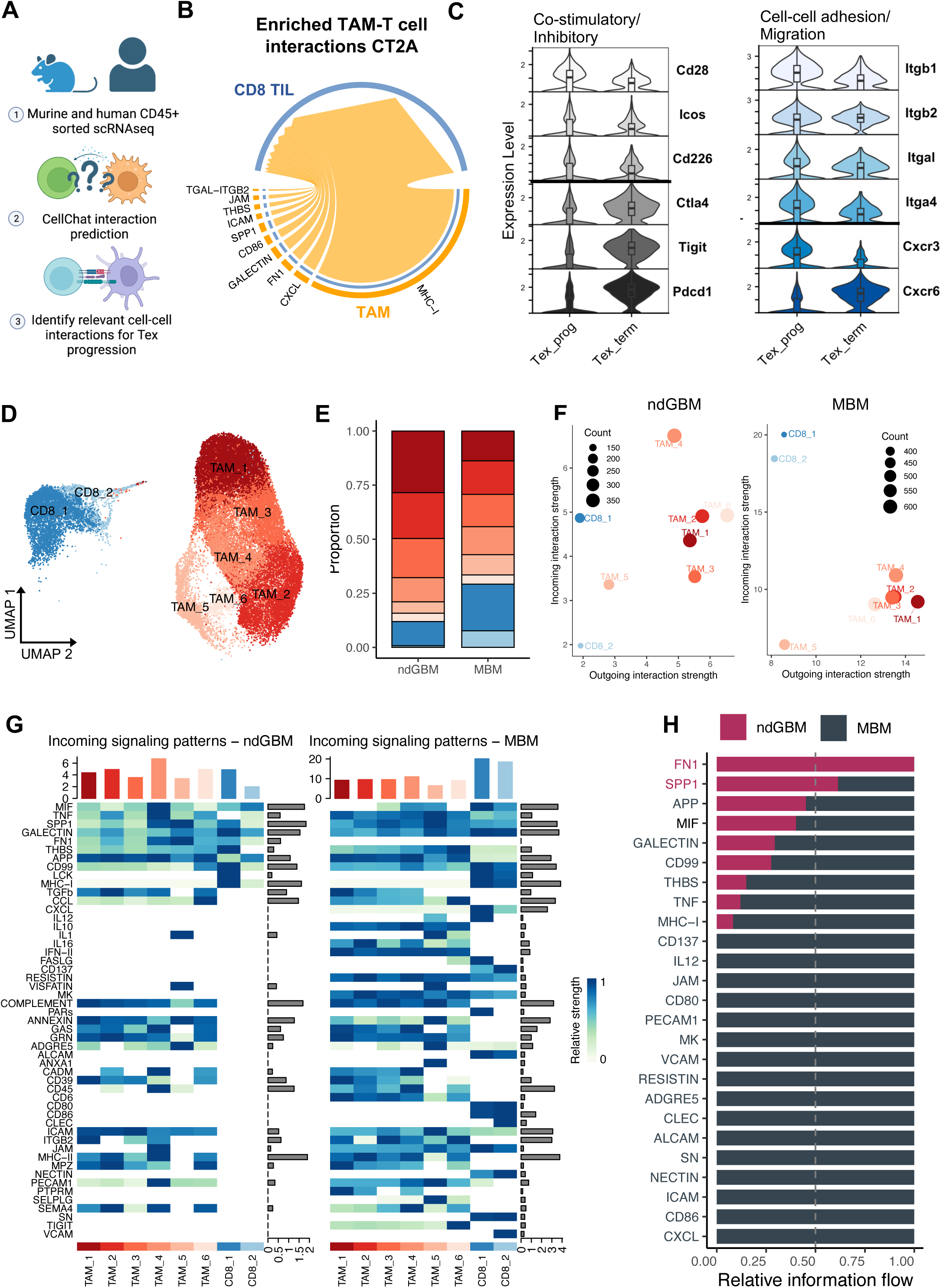
CellChat predicts secondary interactions between TAM and CD8^+^ T cells in mice and humans that may contribute to exhaustion progression. (A) Experimental layout. CD45^+^ sorted scRNAseq from murine GBM, ndGBM and MBM was analyzed with CellChat to determine additional TAM-T cell interactions. Created with Biorender. (B) Enriched predicted interactions with TAM (orange) as the source and CD8 TIL (blue) as the target from murine GBM. (C) Receptor expression in Tex_prog cells vs. Tex_term cells from CD3^+^ sorted dataset from Figure 2 of predicted interactions from 6B. (D) UMAP of human ndGBM and MBM datasets. (E) Frequency of TAM vs. CD8^+^ T cells in ndGBM vs. MBM. (F) Incoming vs. outgoing interaction strength across cell types in ndGBM vs. MBM. (G) Heatmap of all incoming signaling patterns in ndGBM vs. MBM. (H) Significantly enriched predicted pathways in ndGBM (maroon) vs. MBM (grey).

To determine which of the above predicted interactions might also be prevalent in patients, we turned to publicly available human GBM and metastatic melanoma datasets. We integrated 15 newly diagnosed GBM (ndGBM) and 5 treatement-naïve metastatic brain melanoma (MBM) samples and focused on TAM and CD8^+^ T cells for further analysis^39,40^. Once integrated, we derived 8 TAM clusters and 3 CD8 clusters that were shared between tumor types (**Figure 6D**). As expected, the abundance of TAM was greater in ndGBM than in MBM (**Figure 6E**). We then performed cell – cell communication analysis with the human samples using CellChat ^33^. Broadly, CD8^+^ T cells possessed high “incoming interaction” levels but low “outgoing interaction” strength for both tumor types, whereas TAM had comparable incoming/outgoing strengths, highlighting their interconnectedness within the TME (**Figure 6F**). The total interaction strength was much higher in MBM than ndGBM, suggesting the MBM TME, as a whole, exhibits receptor – ligand pairs at a higher level (**Figure 6F**).

Given the higher strength of incoming interactions amongst CD8^+^ T cells, we evaluated which predicted interactions were prevalent between TAM and CD8^+^ T cells across tumor types. Among those predicted, several overlapped with our CellChat analysis of murine GBM, including MHC-I, SPP1, CD86, ICAM, FN1, JAM, CXCL, and GALECTIN (**Figures 6B and 6G**). ndGBM was primarily enriched for cell – cell adhesion interactions, whereas MBM had exhibited a greater representation of inflammatory/migration pathways. These results align with perceptions of GBM as an “immunologically cold” tumor compared to MBM^41,42^. When comparing the intensity of these pathways between tumor types, we again observed an increased strength and number of interactions in the MBM group (**Figure 6H**). The SPP1 and FN1 pathways were increased in the ndGBM samples compared to MBM. TAM SPP1 expression has been previously implicated in furthering levels of T cell exhaustion in brain tumors^43^.

### Increasing PETER via TAM depletion leads to improved αPD1 efficacy in checkpoint blockade-responsive models

Depletion of TAM produced an increase in PETER across multiple tumor models tested (Figures 5A-B and 5K-N). As the maintenance of Tex_prog cells has been shown to be an important consideration in predicting immunotherapy responsiveness^15,18,21,24,29^, we hypothesized that increasing PETER via TAM depletion would likewise enhance the efficacy of αPD1 in tumor models. We chose two tumor models: one that has been traditionally non-responsive to immune checkpoint blockade (CT2A glioma), and one that has been responsive (MC38 colorectal carcinoma). In both models, TAM depletion and αPD1 were administered alone or in combination (**Figure 7A**). In mice with intracranial CT2A-TRP2, TAM depletion was not sufficient to improve survival, or to license a response to αPD1 treatment (data not shown). Analysis of tumor-infiltrating CD8+ T cells showed a trending decrease in the frequency of Tex_term cells in the combo TAM depleted + αPD1 group (**Figure 7B**), corresponding to an increase in PETER (**Figure 7C**). In the MC38 model, however, we indeed observed an enhancement to baseline αPD1 efficacy in the TAM depleted group (**Figure 7D**). There was an increase in PETER among intratumoral T cells in this setting (**Figures 7E and 7F**). These findings suggest that increasing PETER via targeting of TAM has the potential to augment an immune checkpoint blockade response in previously responsive tumors.

**Figure 7:**
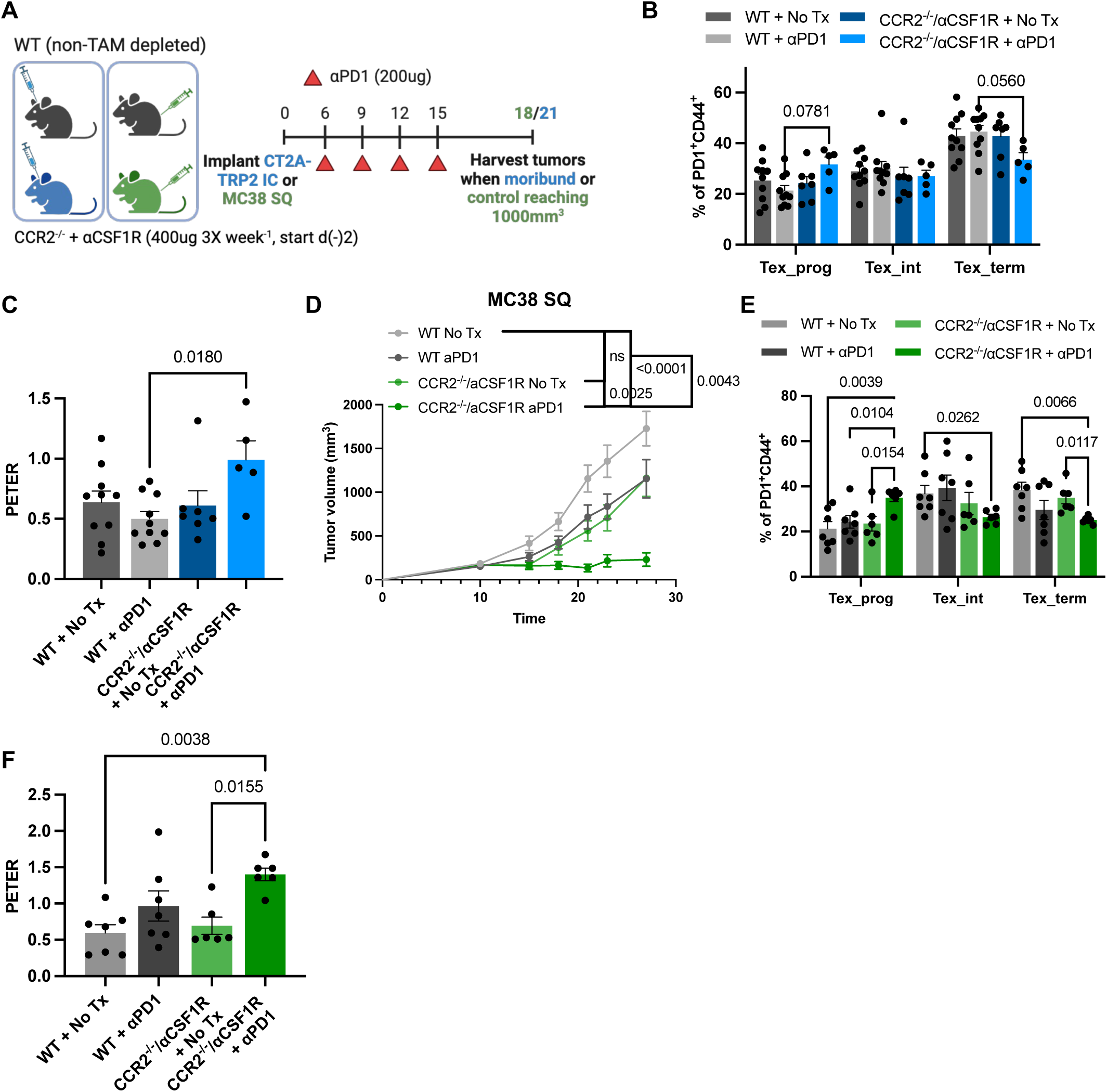
Increasing PETER via TAM depletion leads to improved αPD1 efficacy in checkpoint blockade-responsive models. (A) Experimental layout. TAM and monocytes were depleted in CCR2^−/-^ mice with αCSF1R depletion antibody. αPD1 (200ug) was administered I.P. every 3 days for 4 total injections starting D9 post-injection. 5.0 × 10^4^ CT2A-TRP2 cells were implanted intracranially or 5.0 × 10^5^ MC38 cells were implanted subcutaneously and harvested on D21 or D18, respectively. Created with Biorender. (B, and C) frequency of T cell exhaustion subsets (B) and PETER (C) among mice implanted with intracranial CT2A-TRP2. n = 10 mice/group. (D, E and F) Subcutaneous tumor growth curve (D), frequency of T cell exhaustion subsets (E) and PETER (F) among mice implanted with subcutaneous MC38. n = 10 mice/group. Data are representative of 2 independent experiments. p-values are plotted on individual graphs. Statistical significance was determined by 1-way ANOVA with post-hoc Tukey’s test for multiple comparisons (C and F) or RM-2-way ANOVA with post-hoc Tukey’s test for multiple comparisons (B and E), tumor growth curves were assessed with 2-way ANOVA with post-hoc Tukey’s test. All data are plotted as mean ± SEM.

## DISCUSSION

The abundance of Tex_prog cells within a TME is associated with immunotherapeutic responsivity ^15,20,21,24,42^. Initially identified as the T cell differentiation state remaining reactive to anti-PD1 treatment in chronic infection, Tex_prog is a consistent correlate of ICB response^15,18,23^, and the abundance of Tex_prog cells relative to Tex_term cells can be critical for responsivenss ^24^. Here, we detailed the severe and disproportionate loss of Tex_prog cells over time in GBM and other solid tumors and implicated TAM, but not tumor cells, as the mediators of the Tex_prog to Tex_term transition. Recognizing the importance of this shift from Tex_prog to Tex_term, we, in turn, coined and identify the PETER as an important indicator of the dynamics of T cell exhaustion within the TME. As GBM possesses a particularly low PETER and is likewise unresponsive to ICB, we suggest that a therapeutic goal for licensing improved immunotherapeutic responses may ultimately be finding ways to optimize PETER within a given TME.

Further characterizing Tex_prog cells and Tex_term cells, we observed an enrichment for chemotactic and cell-cell adhesion pathways in Tex_prog cells, and we implicated TAM as driving a subsequent loss of T cell polyfunctionality and stem-like properties as T cells increasingly differentiated toward Tex_term cells in the tumor. In the absence of TAM, we instead observed retention of the Tex_prog population and a resultant increase in PETER, with the retained Tex_prog cells possessing established implications for immunotherapeutic responsivity^16,18,21,23,24^.

The increase in the proportion of Tex_prog cells in the TME in the absence of TAM demonstrates that the impact within the TME is not in the evolution from naïve T cell to Tex_prog cell, but rather at the progression from Tex_prog cell to Tex_term cell (i.e. why we saw an increase in PETER in the absence of TAM). This also suggests the progression from naïve to Tex_prog may occur earlier, i.e. in the tdLN following priming, with secondary antigen presentation on MHC I by TAM in the TME mediating the subsequent progression to Tex_term. Accordingly, we observed only Tex_prog cells within the tdLN, and only at low levels, even at late stages of tumor progression. Even following FTY720 administration, with accompanying retention of Tex_prog cells in the tdLN, no progression to Tex_term cells occurred in the LN. Together, these data support the suggestion that the transition of Tex_prog cells to Tex_term cells occurs in the TME alone. Similar conclusions were reached by another group, who show that partial T cell differentiation to Tex_prog occurs in the LN, but effector (Tex_term) differentiation occurs in the tumor^22^.

Many defining studies for T cell exhaustion were conducted in the chronic infection model of lymphocytic choriomeningitis virus (LCMV)^14,17,44^. In the LCMV model, it is unfortunately impossible to determine whether T cell chronic antigen exposure occurs on the surface of infected target cells or APC, as they are typically one and the same. As a result, a common presumption is that chronic exposure to antigen presented by target cells is a driver of T cell exhaustion progression. In contrast, in tumors, T cell antigen exposure may occur on the surface of tumor cells presenting endogenous antigen, or in the context of TME APC cross-presenting exogenous antigen. Looking to assess whether one antigenic context might be more crucial to the derivation of exhaustion in the TME, we removed the capacity for MHC I presentation from tumor cells and observed no change in the frequency of T cell exhaustion subsets. This suggested that encountering antigen on tumor cells is not required for the differentiation of Tex_prog to Tex_term, leaving intratumoral APC as our next-investiagted source of instigatory antigen-cognate interactions.

In our intial querying of intratumoral APC, DC were a leading logical candidate for exhaustion-eliciting interactions with Tex_prog cells. However, interactions with DC, although present, proved to be neither necessary nor sufficient for the differentiation of Tex_prog to Tex_term. Rather, we found TAM to be a necessary cell type, with TAM MHC I expression especially implicated: Loss of MHC I *ex vivo* on TAM led to decreased accumulation of Tex_term cells, with the degree of Tex_term cell persistence correlating with residual TAM MHC I levels *in vivo*.

Several groups have now highlighted the role of infiltrating myeloid cells, namely TAM and DC, in T cell terminal differentiation. Nixon *et al*. elegantly identify IRF8 as a regulator of increased antigen presentation capacity in infiltrating (and not resident) TAM. When IRF8 is conditionally deleted from TAM using a MafB-iCre X Irf8 floxed model, increased tumor control is observed, and TAM have decreased antigen presentation capacity. The authors identify that TAM have a unique tolerogenic effect on T cells in co-culture that is not observed when T cells are cultured with DC or tumor cells^28^. To add to this body of knowledge, we more directly demonstrated the importance of MHC I, specifically, to the T cell exhaustive process.

Additional TAM – CD8^+^ T cell crosstalk was described by Kersten *et al.* in a murine model of melanoma, whereby CD8-derived CSF1 contributes to TAM recruitment and differentiation, which allows for TAM-T cell interactions^27^. This group also demonstrates a unique capacity for TAM to sustain longer, but more weakly activating interactions with CD8^+^ T cells *in vitro* than DC-CD8^+^ T cell interactions. Likewise, a role for attenuated TCR stimulation is identified in the progression of T cell exhaustion by Lan *et al*^45^. The authors show that suboptimal TCR stimulation accelerates terminal differentiation, whereas robust TCR stimulation reinforces Tex_prog cell self renewal. This is perhaps especially important in TAM-enriched tumors, like GBM and colorectal cancer, since TAM are more likely to provide suboptimal TCR stimulation.

In addition to being the most abundant APC, TAM also demonstrated the greatest propensity for phagocytosis and cross-presentation in the TME. The sheer number of predicted TAM – T cell interactions are important as such interactions are generally only weakly activating and often even immunosuppressive compared to DC – T cell interactions^27,31,46,47^. Our findings, however, highlight a newly appreciated immunoregulatory role for TAM and provide a new focus on determining relevant T cell – TAM interactions for targeting in an effort to better license immunotherapeutic responses.

While antigen cognate interactions with TAM may be a necessary first step in driving TAM – T cell interactions and thus T cell exhaustion within the TME, our data suggest that there are additional secondary interactions that follow and may help facilitate the Tex_prog to Tex_term transition. Other groups proposed co-stimulatory signals through the CD86/CD28 axis as contributors^22^. Our CellChat findings suggested that such interactions may indeed play a role in more inflammatory tumors, such as MBM, but this axis was not prevalent in our human ndGBM dataset. Interestingly, we identified SPP1, which binds to integrins (CD44 or CD29) on CD8^+^ T cells, as a highly predicted component of interaction between TAM and CD8^+^ T cells in in both murine and human tumors. Importantly, SPP1 has a T cell exhaustive role in the setting of GBM. Kilian *et al.* identify a pathway in which the absence of CD4^+^ T cell – TAM MHC II interactions leads to increased secretion of SPP1 by TAM, which suppresses surrounding CD8^+^ T cells and increases their expression of exhaustion-associated factors^43^. Further studies are ultimately needed to better address the nature and role of specific interactions between TAM and CD8^+^ T cells.

Given the known importance of Tex_prog cells in mediating the response to immunotherapy, we hypothesized that an increase in the PETER would yield enhanced immunotherapeutic responses for a given tumor. Importantly, we observed this to be true in a tumor previously responsive to checkpoint blockade (MC38), but were unable to newly license a response in tumors that were previously entirely checkpoint-resistant (CT2A). This finding may reflect some of the additional hurdles to immunotherapeutic success that have been historically present with GBM.

Taken together, we have highlighted TAM – T cell interactions and TAM antigen presentation as critical for driving CD8^+^ T cell exhaustion progression in GBM and other solid tumors. Future work will focus on defining additional specific TAM – T cell interactions and their relative contributions to the differentiation of Tex_prog to Tex_term. The goal remains to promote a TME with a heightened PETER and renewable Tex_prog populations in an effort to better license immunotherapeutic responses.

## ACKNOWLEDGMENTS

Dr. Fecci is funded by a Cancer Research Institute (CRI) Arash Ferdowsi Lloyd J. Old STAR Award. We would like to thank the Duke Molecular Genomics Core, Duke Human Vaccine Institute Flow Cytometry Core, and the Duke Cancer Institute Flow Cytometry Core.

## AUTHOR CONTRIBUTIONS

Conceptualization, J.WP and P.E.F.; Methodology, J.WP., W.H.T., A.M.H-M., L.P.W., D.S.W., E.L., and K.W.; Investigation, J.WP.; Resources, J.WP., W.H.T., A.M.H-M., and J.B.F.; Writing – Original Draft, J.WP.; Writing – Review & Editing, J.WP., A.M.H-M., W.H.T., L.P.W., S.J.L. and P.E.F.; Supervision, P.E.F.; Funding Acquisition, J.WP., K.A., and P.E.F.

## STAR METHODS

### RESOURCE AVAILABILITY

#### Lead contact

Further information and requests should be directed to Peter Fecci.

#### Materials availability

Materials generated by this study are available upon request.

#### Data and code availability

Raw FASTQ files and associated single cell RNA and TCR sequencing processed files have been uploaded to the NCBI Gene Expression Omnibus (GEO) database under GSE236243 (raw and processed data files are available upon request). Custom R scripts for TCR, and RNA analysis are also available upon request.

### EXPERIMENTAL MODEL AND SUBJECT DETAILS

#### Mice

All mice were used in accordance with National Institutes of Health and the Duke University Institutional Animal Care and Use Committee guidelines. C57BL/6J (cat #: 664), CCR2^−/-^ (cat #: 4999), CD11c-DTR (cat #: 4509), CD45.1 (cat #: 2014), and β2m^−/-^ (cat #: 2087) were purchased from Jackson Laboratory at 7-8 weeks.

#### Tumor cell lines

C57BL/6 syngeneic CT2A glioma was provided by Robert L. Martuza (Massachusetts General Hospital)^48^. B16F0 melanoma, E0771 breast cancer, and Lewis lung cancer (LLC) were purchased from ATCC. MC38 was purchased from Kerafast. CT2A-TRP2 was generated by overexpressing a TRP2-containing lentiviral plasmid. CT2A-TRP2-OVA was generated through lentiviral transduction of parental CT2A-TRP2 with an OVA-containing lentiviral plasmid. CT2A-GFP-LUC was kindly provided by Alexa Bramall and Nalini Mehta (Duke University)^49^. CT2A-TRP2-β2m^−/-^ was made via CRISPR/Cas9. sgRNA targeting murine β2m (gRNA sequence: AGUCGUCAGCAUGGCUCGCU) and Cas9 nuclease were purchased from STEMCELL technologies (Cat#: 200-0013, 76002, respectively). sgRNA (5.8 µg) was incubated at room temperature with 10.8 µg Cas9 protein (STEMCELL Technologies, Cat#: 76002) in Opti-MEM™ + GlutaMAX™ (ThermoFisher Scientific Cat#: 51985034) for 20 min prior to electroporation. sgRNA/Cas9 was then added to 2.5 × 10^5^ CT2A-TRP2 cells in Opti-MEM™ + GlutaMAX™. Cells were electroporated in a 0.2 cm cuvette using a BioRad Gene Pulser XCell™ in 100ul total volume (exponential decay, 1000 uF, 155 V). Cells were FACS isolated after 96 h on H-2K^b^/H-2D^b^ negative cells. CT2A-TRP2-β2m^−/-^ cell line was established from a single cell clone, cultured as described below. CT2A, CT2A-TRP2, CT2A-TRP2-OVA, CT2A-TRP2-β2m^−/-^, CT2A-GFP, E0771, LLC, and MC38 were cultured in complete DMEM (Gibco 11995-065, 10% FBS). B16F0 was cultured in complete DMEM + 1mM L-glutamine. All cell lines were authenticated and tested negative for mycoplasma, and interspecies contamination by IDEXX Laboratories.

### METHOD DETAILS

#### Tumor cell line preparation and injection

All cell lines were cultured at 37°C in 5% CO_2_, in their respective media. For tumor inoculation, all cell lines were taken out of culture using 0.05% Trypsin, washed once with media and once with PBS. Cells were resuspended in PBS for subcutaneous injections (200 µl injection volume) and a 1:1 mix of PBS and methylcellulose for intracranial injections (5 µl injection volume). 1 10^4^ CT2A, 5 10^4^ CT2A-TRP2, 5 10^4^ CT2A-TRP2-β2m^−/-^, 5 10^4^ CT2A-TRP2-OVA, 2.5 10^4^ CT2A-GFP-LUC, 1 10^3^ B16F0, 1 10^3^ E0771, and 1 10^3^ LLC were injected intracranially. 5 10^5^ CT2A, 2.5 10^5^ E0771, 1 10^5^ B16F0, 5 10^5^ MC38, were injected subcutaneously. Mice were monitored and sacrificed in accordance with Duke University Institutional Animal Care and Use Committee guidelines.

#### In vivo antibody administration (αCD8, αCSF1R, αPD1)

For macrophage depletion experiments, 400µg αCSF1R (AFS98, BioXCell) was administered 3 times weekly starting 2 days before tumor implantation into CCR2^−/-^ mice, as described previously^36^.

For CD8^+^ T cell depletion, mice received 200ug anti-CD8α (clone 2.43; BioXCell) daily for 3 days prior to tumor implantation, then every 5 days.

For αPD1 treatment, mice received 200ug αPD1 (clone RMP1-14; BioXCell) every 3 days starting on D6 for 4 total treatments.

#### Dendritic cell depletion

CD11c-DTR or control C57Bl/6J mice were injected intraperitoneally with 5ng/g diptheria toxin (Sigma-Aldrich, D0643) every 3 days, starting D8 after tumor implantation for 4 treatments as previously described^2^.

#### Microglia depletion

C57Bl/6J mice were put on PLX3397 diet (290 mg/kg) or a control diet formulated in AIN-76A standard chow as previously described^50^.

#### Mouse brain tumor tissue processing

For processing intracranial tumors, brains were harvested as before^32^. In brief, mice were transcardially perfused with 10ml PBS + 1% heparin. The tumor-bearing hemisphere was mechanically dissociated with a dounce tissure homgenizer, then enzymatically dissociated (0.05mg ml^−1^ Liberase DL (Roche), 0.05 mg ml^−1^ Liberase TL (Roche), 0.2 mg ml^−1^ Dnase I (Roche) in HBSS with Ca^2+^ and Mg^2+^) for 15 min at 37 °C. The single cell suspension was then washed with PBS and resuspended in RBC lysis buffer (Invitrogen) for 3 min. After quenching the lysis reaction with PBS, the cell suspension was passed through a 70 µm filter. Myelin was removed from the sample with Percoll density gradient centrifugation. Samples were resuspended in 30% Percoll (Sigma-Aldrich) and centrifuged at 500 g for 20 min at 18 °C with low brake. The myelin layer was aspirated, and cells were washed with PBS and counted for subsequent flow cytometry staining.

#### Flow cytometry

For cytokine re-stimulation, samples were resuspended in RPMI + 10% FBS at 0.5 - 1 10^7^ cells/ml in a 48 well plate after being counted. PMA (50 ng/ml) + Ionomycin (1000 ng/ml) + GolgiStop™ (BD Biosciences) + GolgiPlug™ (BD Biosciences) were added and then samples were incubated for 4 h at 37 °C.

If no re-stimulation was necessary, samples were resuspended at 1-2 10^7^ cells/ml in 100µl PBS and transferred to a 96-well plate. Cells were resuspended in Zombie Aqua Viability Dye (1:500, Biolegend) and incubated for 25 min at RT.

For extracellular staining, samples were resuspended in 50 µl TruStain FcX Plus (1:500, Biolegend) in FACS (PBS + 2% FBS) for 15 min at RT. After blocking, 50 µl of a 2X antibody mastermix were added and samples were incubated for 25 min at 4 °C. Stained samples were washed with FACS, then fixed with 2% formaldehyde in PBS for 20 min at 4 °C.

For intracellular staining, cells were instead fixed and permeabilized with a 1X FoxP3 Fixation/Perm buffer (eBioscience FoxP3/Transcription Factor Staining Buffer Set, Thermo Fisher Scientific) for 30 min at 4 °C. Intracellular antigens were stained for 30 min in 1X perm buffer at RT.

Before sample acquisition, 10 µl of Count Bright counting beads (Thermo Fisher Scientific) were added to each sample. Samples were acquired on an LSRII or Fortessa (BD Biosciences) using FACS Diva software v.9 and analyzed using FlowJo v.10 (Tree Star).

#### Immunofluorescence

Brains from PLX3397 treated tumor-bearing mice were harvested after intracardial perfusion with 10ml 1% PFA and 10ml PBS. Brains were left in 1% PFA for 6 h, then washed and placed in 10% sucrose for 48h. Once brains had sunk in the solution, they were washed and snap frozen in tissue freezing media (General Data Company) then kept at -80 °C. Frozen brains were sectioned at 25 µm and mounted onto slides. Sections were rehydrated with PBS, then permeabilized with Triton-X and washed with PBS. Permeabilzed sections were blocked using PBS and 2% BSA. Primary and secondary antibodies were subsequently stained in PBS for 1 h at RT each. Slides were imaged on a Leica STED confocal microscope (Leica). FIJI was used to process and analyze images.

#### *Ex vivo* TRP2-TCR transgenic T cell generation and adoptive transfer

Spleens were harvested from CD45.1 mice. Spleens were processed through 70µm filters then red blood cells were lysed with RBC lysis (Invitrogen). Splenocytes were plated at 2 10^6^ cells/ml in RPMI + 10% FBS, 50 U/ml IL-2, 2.5 µg/ml Concavalin A, L-glutamine (Gibco), Pen/Strep (Gibco), 50 µM βME and kept in culture for 48h. Cells were then retrovirally transduced with the pMX-TRP2-TCRβ-2A-α vector (a kind gift from Dr. Schumacher, Netherlands Cancer Institute) as described previously^51^. This TCR recognizes the TRP2_180-188_ epitope in the context of H-2K^b^. Briefly, HEK293T cells were transfected with pCL-Eco and the TRP2 vector using Lipofectamine 2000 (Invitrogen). Transduction was then performed in non-tissue culture treated 24-well plates, previously coated with Retronectin (25 µg/ml) (Takara) by combining viral supernatant and activated splenocytes and spinning for 1.5 h at 2000 RPM. Transgenic (Tg) TRP2-TCR T cells were cultured for another 3 days before use, splitting daily. 10 × 10^6^ or 2.5 × 10^5^ Tg TRP2-TCR T cells were adoptively transferred through tail vein or intracranial injection in 100 µl PBS or 24 µl PBS, respectively.

OT1 T cells were isolated from OT1 mice by culturing OT1 splenocytes as above for 48 h.

#### Chimera Generation

Bone marrow was harvested from 7 week old donor mice (either C57BL/6J or β2m^−/-^). Recipient mice (CD45.1) received whole-body irradiation with a 9 Gy dose from a Cesium irradiator (Mark I-68A ^137^Cs irradiator, JL Shepherd and Associates). Recipient mice then received an intravenous infusion of 10 × 10^6^ donor bone marrow cells in 100 µl PBS. Recipient mice were then put on antibiotic-treated water for 2 weeks post bone marrow transfer.

After 10 weeks of reconstitution, chimerism was validated in the blood, with >95% chimerism in all queried mice. Then, CT2A-TRP2 or CT2A-TRP2-β2m^−/-^ was implanted intracranially into chimeras. On D7 post-implantation, 10 × 10^6^ TRP2-TCR T cells were adoptively transferred intravenously.

#### *In vitro* co-culture

OT1 T cells were isolated from OT1 transgenic mice and activated as previously described by Wu et al.^38^. Briefly, OT1 T cells were isolated using negative CD8^+^ T cell bead isolation (Stemcell), then co-cultured with OVA-peptide-pulsed splenic CD11c^+^ DC for 48h. After activation, OT1 were washed then co-cultured with OVA-pulsed TAM and tumor cells (CT2A) at an E:T of 3:1 for 24h in T cell media + 10U/mL IL2 (peprotech). TAM were isolated from IC CT2A tumors 20 days post-injection, where they were sorted as CD45^+^CD11b^+^F4-80^+^.

#### scRNAseq library generation

CD3^+^CD4^+^ or CD3^+^CD8^+^ were sorted from D13 (n = 16) and D19 (n = 4) intracranial tumors. Four mice were pooled for each D13 replicate to achieve desired number of cells. 10,000 total cells from each timepoint were targeted with an average read depth of 36,000 reads per cell for the gene expression library. Reads were sequenced on a NovaSeq S1 flow cell. For the TCR library, 9,000 reads per cell were targeted and sequenced on a NovaSeq SP.

##### 10x Transcriptome library prep

GEMs are generated by combining barcoded Single Cell 5’ Gel Beads, Master Mix, Partitioning Oil and single cells on a Chromium microfluidic Chip. Immediately following GEM generation, the Gel Bead is dissolved and any co-partitioned cell is lysed. Oligonucleotides containing an Illumina R1 sequence (read 1 sequencing primer), a 16 nt 10x Barcode, a 10 nt unique molecular identifier (UMI), and 13 nt template switch oligo (TSO) are released and mixed with the cell lysate and a Master Mix containing reverse transcription (RT) reagents and poly(dT) RT primers. Incubation of the GEMs produces 10x Barcoded, full-length cDNA from polyadenylated mRNA.

GEMs are then broken and pooled and the 10x Barcoded first-strand cDNA is purified from the post GEM-RT reaction mixture and amplified. Full-length V(D)J segments (10x Barcoded) are enriched from amplified cDNA via PCR amplification with universal primer for 5’ adapter and successive nested primers specific to the TCR and/or BCR constant regions. Enrichment and library preparation of the TCR and BCR constant regions takes place in two separate workflows using the same amplified cDNA stock.

Enzymatic fragmentation and size selection are used to generate variable length fragments that collectively span the V(D)J segments of the enriched TCR and/or BCR transcripts prior to library construction. P5, P7, a dual sample index, and an Illumina R2 sequence (read 2 primer sequence) are added via End Repair, A-tailing, Adaptor Ligation, and Sample Index PCR. The final libraries contain the P5 and P7 priming sites used in Illumina sequencing. Sequence is generated using Illumina short read paired end sequencing.

#### scRNAseq data processing and analysis

The Cell Ranger pipeline (10X Genomics) was used to perform sample demultiplexing and to generate FASTQ files for the gene expression and ADT libraries. FASTQ files were demultiplexed from the raw sequencing reads (bcl2fastq, v2.20), aligned to the mouse mm10 reference genome (cellranger, v4.0.0), and raw gene count matrices were generated using STAR (v2.7.5c).

Downstream analysis was performed using the R software Seurat package^52^ (v4.0.3, http://satijalab.org/seurat/). Hashtag oligos, which corresponded to biological replicates, were demultiplexed using the HTODemux function and appended to the meta-data for each sample. Low-quality cells, expressing less than 200 genes, and genes expressed by fewer than three cells were removed. The gene expression matrix for each genotype was then concatenated using the merge function in Seurat. The percentage of mitochondrial gene content was calculated using the Mouse.MitoCarta3.0^53^ gene set and the PercentageFeatureSet function in Seurat. The Seurat object was converted to a SingleCellExperiment object, and outlier exclusion was performed in scater (v.1.18.6). Using the isOutlier function in scater, cells were discarded if their percentage of mitochondrial gene content, number of expressed genes, or number of reads for a given cell was considered an outlier. A total of 6,742 CD8^+^ cells passed QC and the SingleCellExperiment object was then converted back into a Seurat object. Normalization and regression of cell cycle scoring and percent mitochondrial gene content were performed using the SCTransform function in Seurat. Principal component analysis (PCA) was performed on all genes, and the number of principal components to be utilized in further analysis was determined heuristically using the elbow method. Eleven principal components were used for clustering and dimensionality reduction using FindNeighbors, RunUMAP, and a resolution of 0.5 was used for FindClusters in Seurat. This approach identified six distinct cell clusters. All subsequent analyses were conducted with this cleaned object.

For TCRSeq analysis, scRepertoire standard workflow was utilized^54^. In brief filtered contig outputs from the 10X Genomics Cell Ranger Pipeline were loaded into R (v4.1.2) and combined into a list using combineTCR. Next combineExpression was used to attach clonotypic information to the Seurat object. Visualizations were made using the scRepertoire vignette.

For exhaustion signature score generation, the built in AddModuleScore function from Seurat was utilized with a gene list from Im et al. 2016^18^. A detailed list can be found in Table S1. For pseudotime analysis, the object was passed into Monocle v3 without any changes to the UMAP dimensional reduction.

#### Publicly available scRNA-seq data analysis

Human newly diagnosed GBM and metastatic brain melanoma datasets were downloaded from the GEO (accession # :GSE160189, GSE200218). Datasets were QC’d as above, removing cells with high mitochondrial percentage and those that lacked expression of PTPRC1 (non-immune cells). Then datasets were integrated using the standard Seurat protocol^55^.

#### Cellchat Analysis Mouse/Human

For murine CellChat analysis utilizing the CD45^+^ sorted dataset generated by Tomaszewski et al^32^, we subsetted the object to keep CD8 TIL, Apoe+ TAM, Nos2+ TAM, DC1, DC2, DC3, pDC and microglia from the WT sample. We then fed this cleaned object into the CellChat pipeline with built in murine ligand-receptor database. Following the recommended vignette, we visualized the enrichment of the MHC-I pathway across cell types. Additionaly we visualized the strength and number of interactions with CD8 TIL as the target cells.

Human CellChat analysis followed a similar workflow, except the human ligand-receptor database was utilized. Additionally, integrated ndGBM and MBM was subsetted so samples of the same tumor type were analyzed separately, as suggested by the CellChat vignette.

### QUANTIFICATION AND STATISTICAL ANALYSIS

#### Statistical Analysis

Unless specifically noted, all data are representative of ≥2 separate experiments. Experimental group assignment was determined by random designation. Statistical analyses were performed using GraphPad Prism software. Error bars represent ± Standard Error of the mean (SEM) calculated using Prism. Specific statistical tests used were paired t-tests, unpaired t-tests, and one/two-way ANOVA unless otherwise noted. P-values <0.05 were considered statistically significant. Investigators were not blinded to group assignment during experimental procedures or analysis.

---

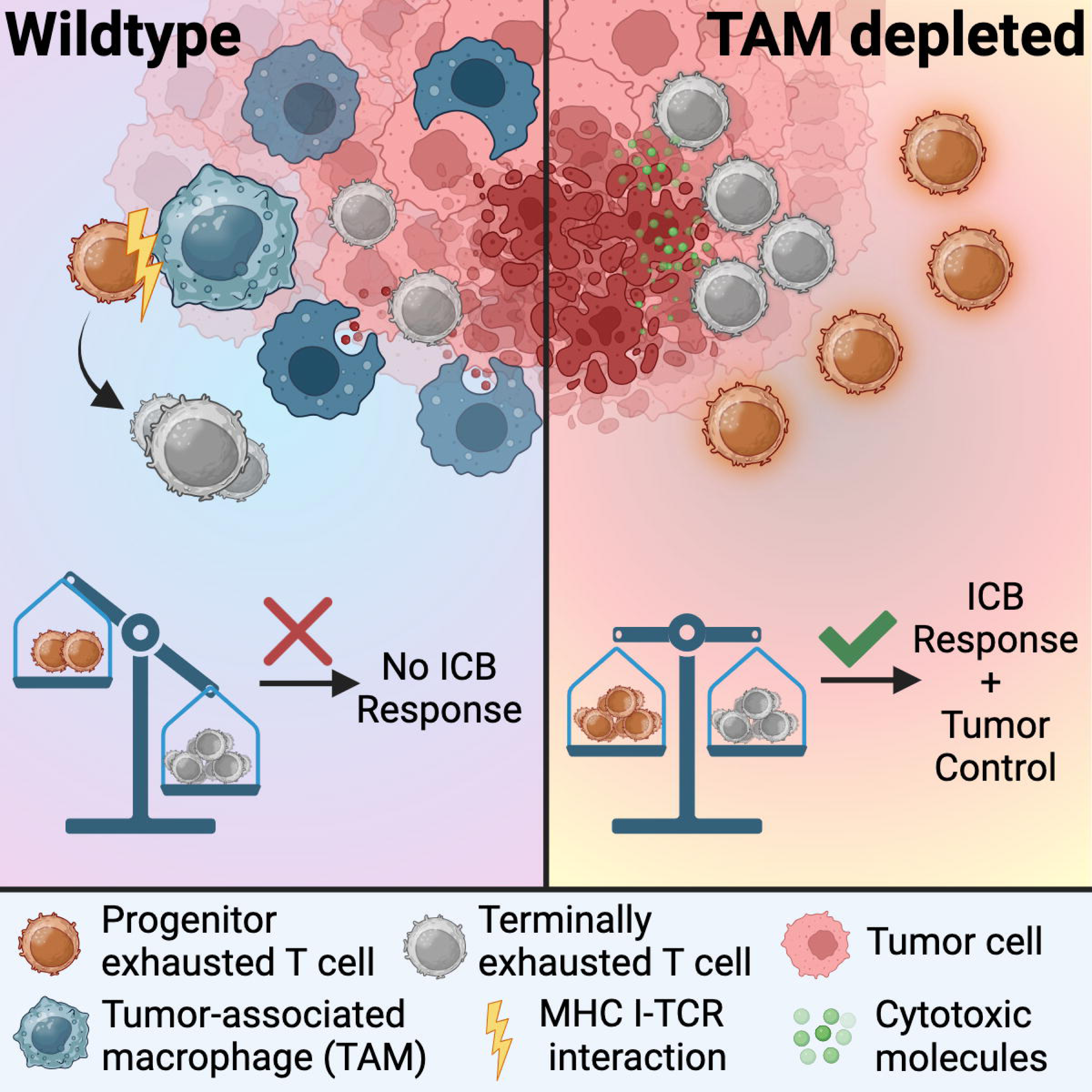

## Supplemental Figures

**Figure S1:**
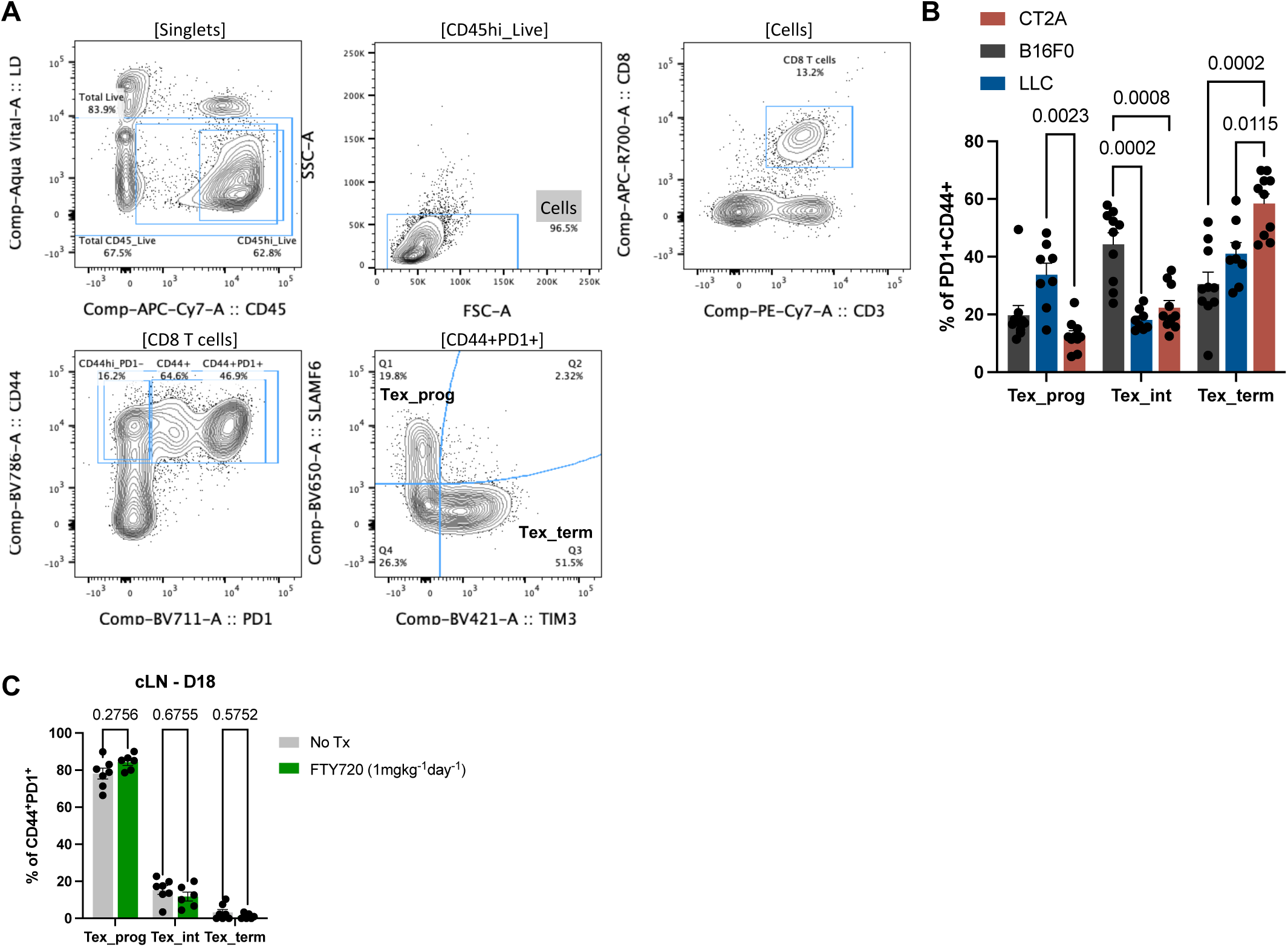
Phenotyping T cell exhaustion subsets in tumor and draining lymph node of various solid tumor models. (A) Gating strategy for Tex_prog and Tex_term. (B) Frequency of Tex subsets of PD1+CD44+ CD8+ T cells in intracranial B16F0, LLC or CT2A. n = 8-10 mice/group. (C) Mice were implanted with 10×10^3^ CT2A tumor cells intracranially and treated IP with 1mg/kg FTY720 daily on days 8-18. Cervical lymph nodes were harvested when mice were moribund (D18). n = 6-7 mice/group. All of the data are means ± SEMs; p-values are shown on individual plots as determined by 1-way ANOVA and post-hoc Tukey test (B and C).

**Figure S2:**
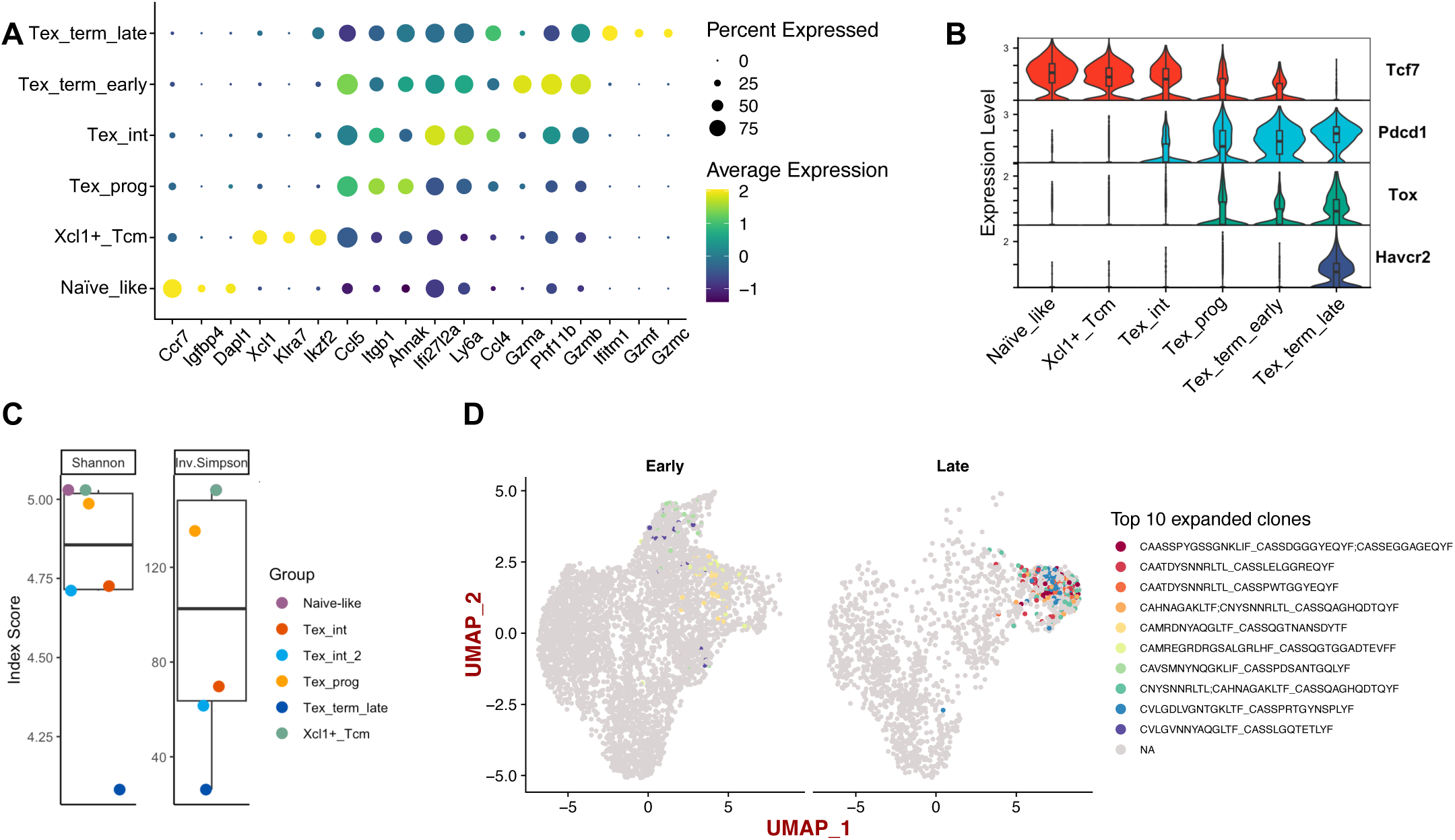
Transcriptional characterization of CD8^+^ T cells in CT2A glioma. (A) Dot plot of top 3 genes for each cluster. (B) Violin plot of *Tcf7, Pdcd1, Tox,* and *Havcr2* expression across clusters. (C) Shannon and Inverse Simpson diversity indices across clusters. (D) 10 most expanded TCR clones across timepoints are concentrated to Tex_int and Tex_term clusters.

**Figure S3:**
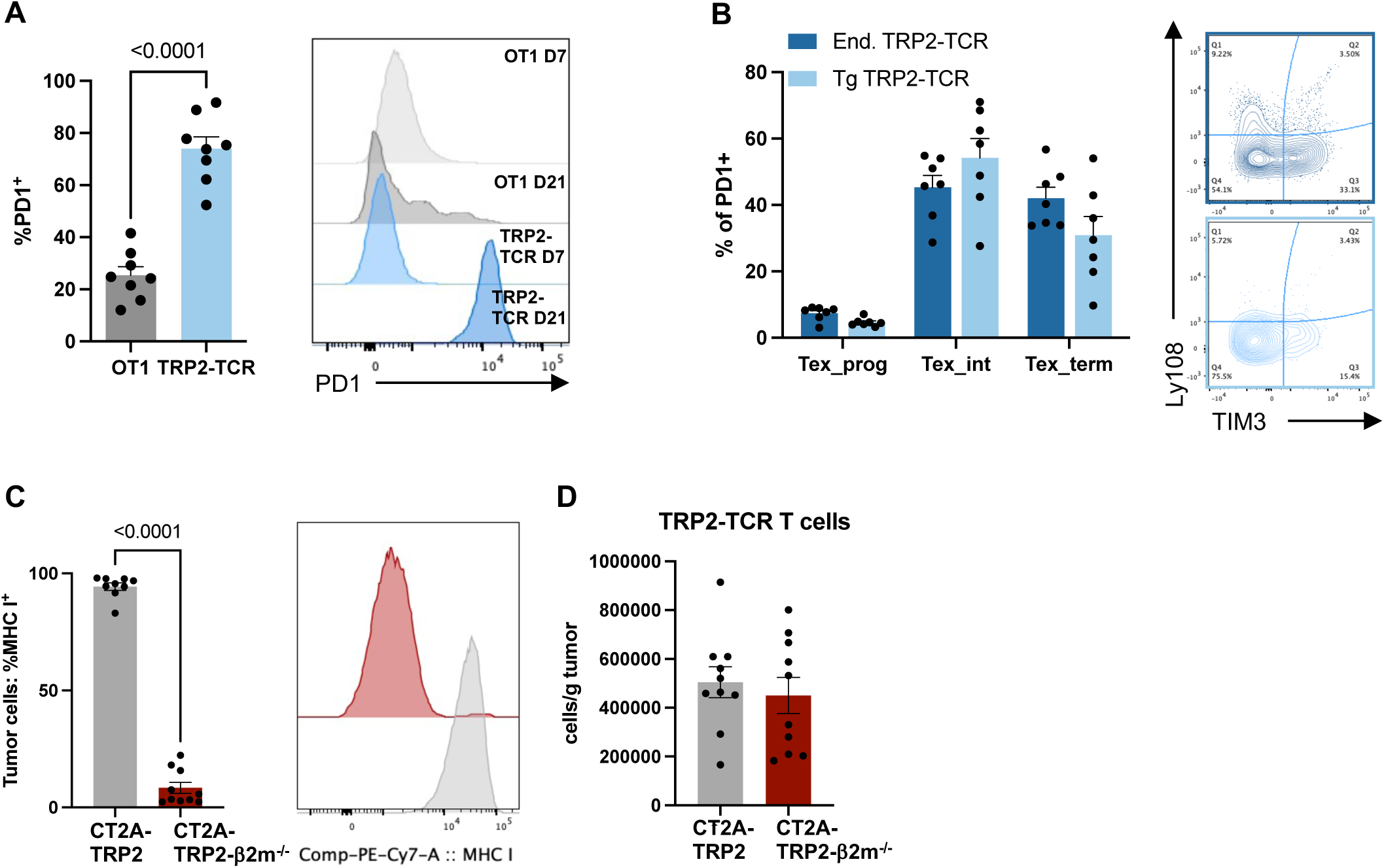
Antigen-specific stimulation is required for T cell differentiation in the tumor. (A) PD1 expression on transferred OT1 vs. TRP2-TCR Tg T cells on day of adoptive transfer (D7) and day of tumor harvest (D21) and representative histogram. n = 7 mice/group. (B) Frequency of Tex subsets of PD1+ endogenous or transgenic TRP2-TCR T cells and representative contour plot. n = 7 mice. (C) gMFI of MHC I in CT2A-TRP2 vs. CT2A-TRP2-B2mKO validating knockdown of MHC I. n = 10 mice/group. (D) Number of TRP2-TCR T cells/g of tumor in CT2A-TRP2 vs. CT2A-TRP2-B2mKO. n = 10 mice/group. p-values are plotted on individual graphs. Statistical significance was determined by two-sided unpaired t-test (A, C, D) or 2-way RM ANOVA with post-hoc Tukey’s test for multiple comparisons (B). All data are plotted as mean ± SEM.

**Figure S4:**
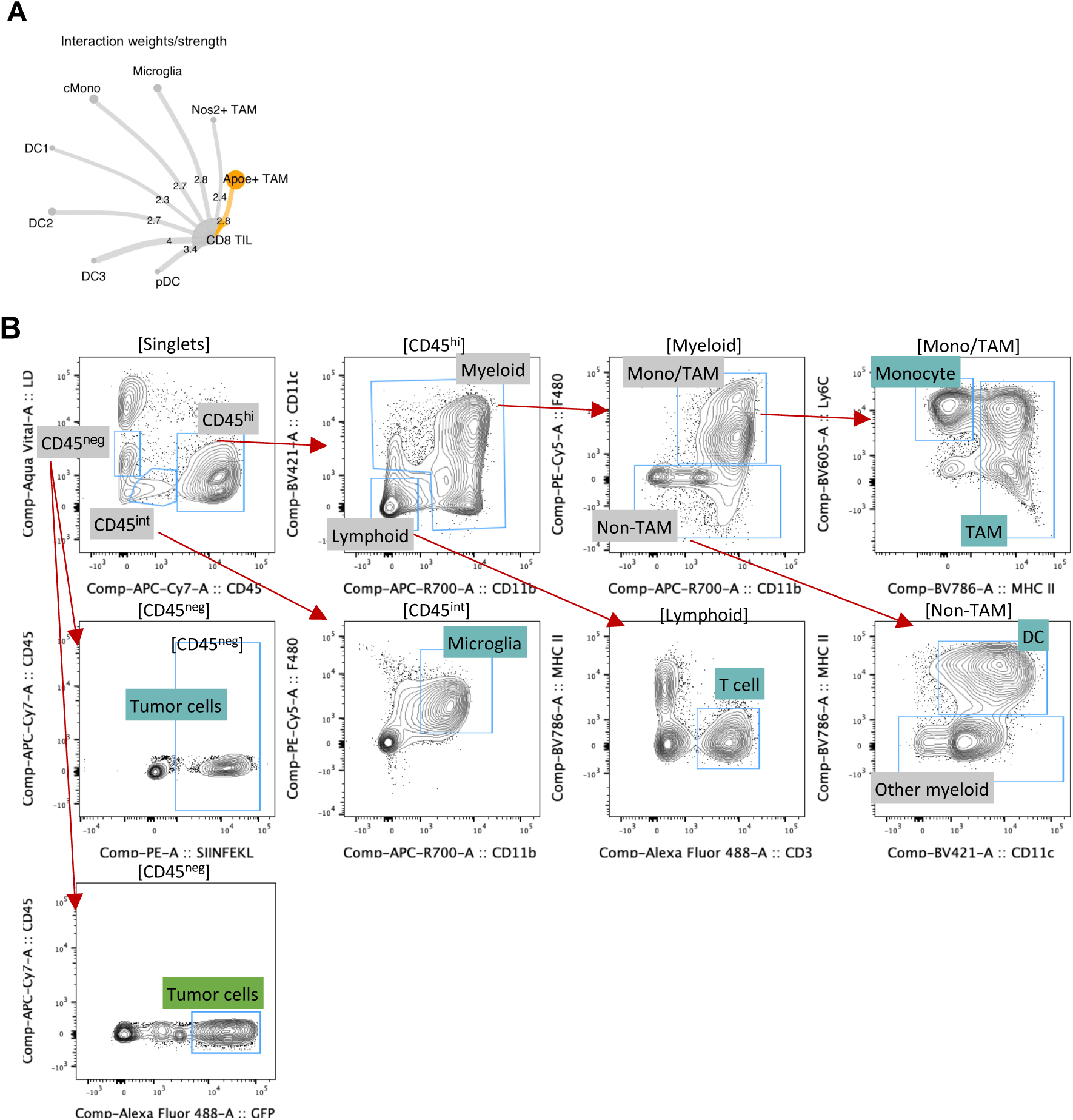
Investigating APC as critical mediators of T cell exhaustion progression in the tumor. (A) Interaction strength across APC with CD8 TIL as the target. (B) Gating strategy for identification of intratumoral APC.

**Figure S5:**
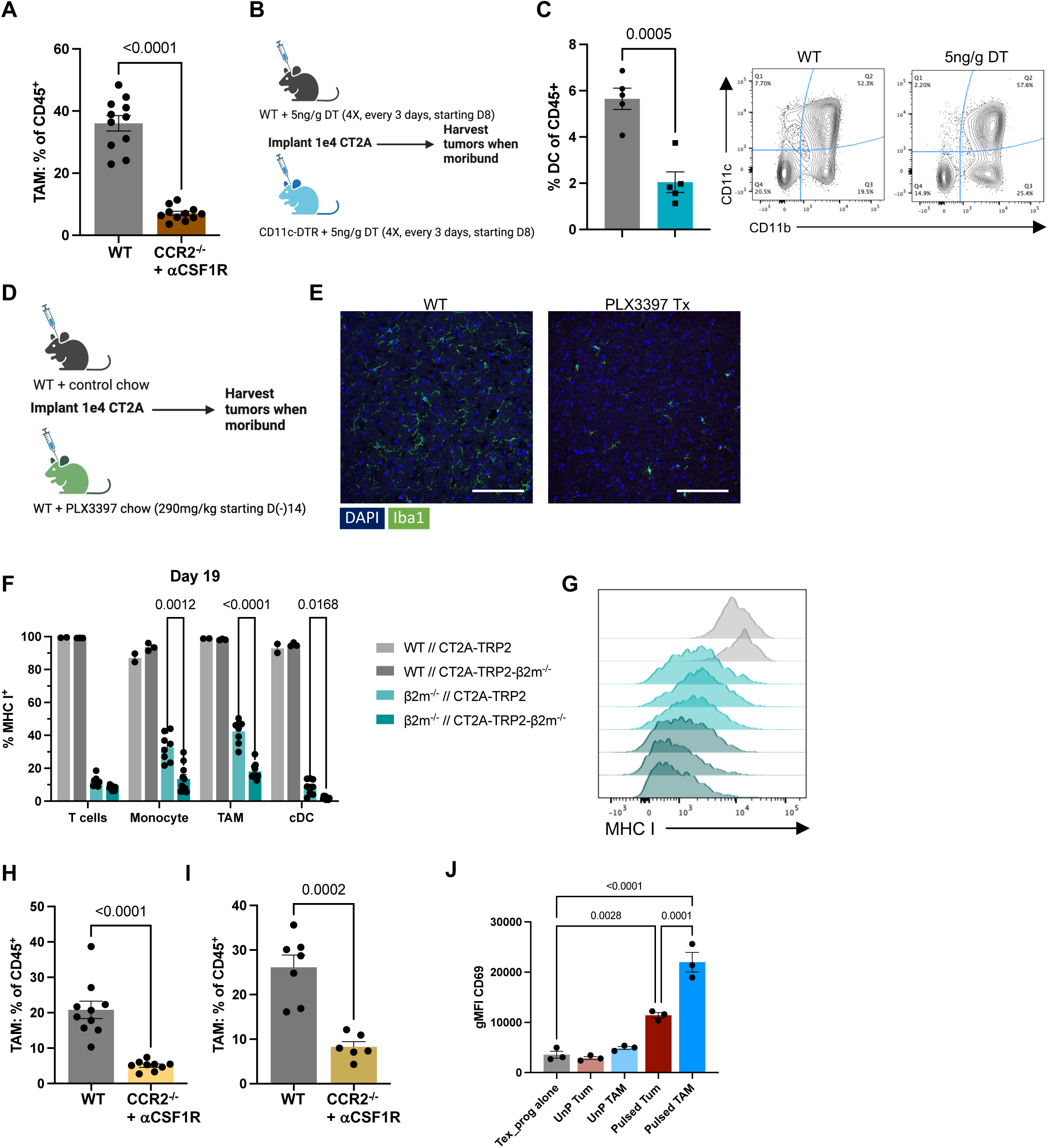
Depletion of TAM, DC and microglia from glioma and melanoma tumor models. (A) Frequency of TAM of CD45^+^ cells. Data pooled from 2 independent experiments. n = 11 mice/group. (B) Experimental layout. DC were depleted in CD11c-DTR mice with 5ng/g DT, every 3 days for 4 total treatments, starting D8. Created with Biorender.com. (C) Frequency of CD11c+CD11b-DC in WT vs. CD11c-DTR mice. (D) Experimental layout. Microglia were depleted in WT mice with 290mg/kg PLX3397 chow. Chow was provided starting D(-)14. Created with Biorender.com. (E) Representative immunofluorescence image of WT vs. PLX3397 treated non-tumor brain. Scale bar = 100um. (F) Chimera validation. Frequency of MHC I+ in WT è WT (WT) or β2mKO è WT (β2mKO) chimeras across T cells, TAM and cDC. n = 8-10 mice/group. (G) Representative histogram of MHC I expression on TAM in the setting of CT2A-TRP2 (Teal) vs. CT2A-TRP2-B2mKO (Dark Teal). (H) Frequency of TAM of CD45^+^ cells in B16F0 IC. n = 9-10 mice/group. (I) Frequency of TAM of CD45+ cells in B16F0 SQ. Data pooled from 2 independent experiments. n = 6-7 mice/group. (J) Expression of CD69 on OT1 T cells when co-cultured with TAM or tumor. n = 3 biological replicates. Data are representative of 2 independent experiments. p-values are plotted on individual graphs. Statistical significance was determined by two-sided unpaired t test (A, H, I), 1-way ANOVA with post-hoc Tukey’s test or RM-2-way ANOVA with post-hoc Tukey’s test for multiple comparisons (F). All data are plotted as mean ± SEM.

