## Supplemental Figures and supp table for "Antigen presentation by tumor-associated macrophages drives tumor-infiltrating T cells from a progenitor exhaustion state to terminal exhaustion"

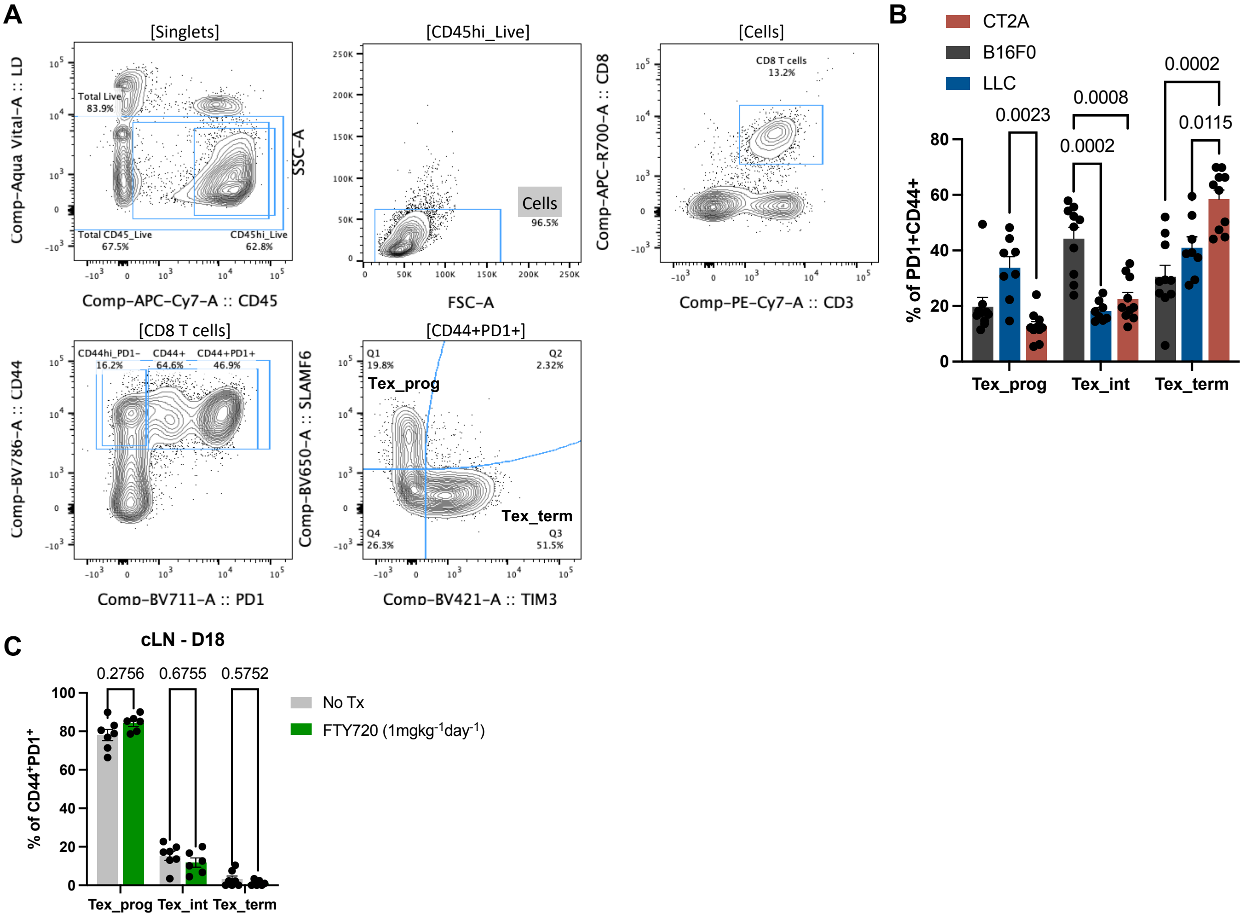


**Figure S1: Phenotyping T cell exhaustion subsets in tumor and draining lymph node of various solid tumor models, related to Figure 1.**
(A) Gating strategy for Tex_prog cells and Tex_term cells. (B) Frequency of Tex subsets of PD1+CD44+ CD8+ T cells in intracranial B16F0, LLC or CT2A. n = 8-10 mice/group. (C) Mice were implanted with $10x10^3$ CT2A tumor cells intracranially and treated IP with 1mg/kg FTY720 daily on days 8-18. Cervical lymph nodes were harvested when mice were moribund (D18). n = 6-7 mice/group. All of the data are means ± SEMs; p-values are shown on individual plots as determined by 1-way ANOVA and post-hoc Tukey test (B and C).

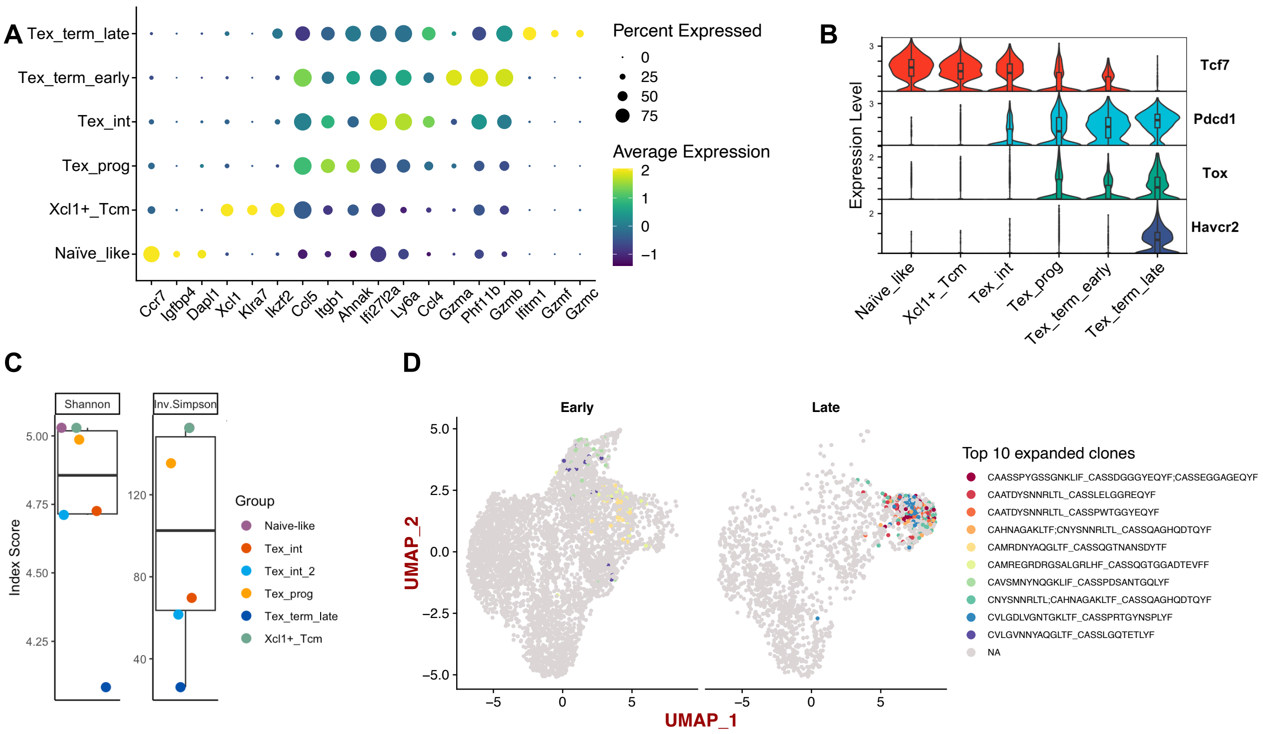


**Figure S2: Transcriptional characterization of CD8⁺ T cells in CT2A glioma, related to Figure 2.** (A) Dot plot of top 3 genes for each cluster. (B) Violin plot of Tcf7, Pdcd1, Tox, and Havcr2 expression across clusters. (C) UMAP of cloneType across biological replicates. n= 4 per timepoint. (D) Shannon and Inverse Simpson diversity indices across clusters, calculated with the scRepertoire package. (E) 10 most expanded TCR clones across timepoints are concentrated to Tex_int and Tex_term clusters.

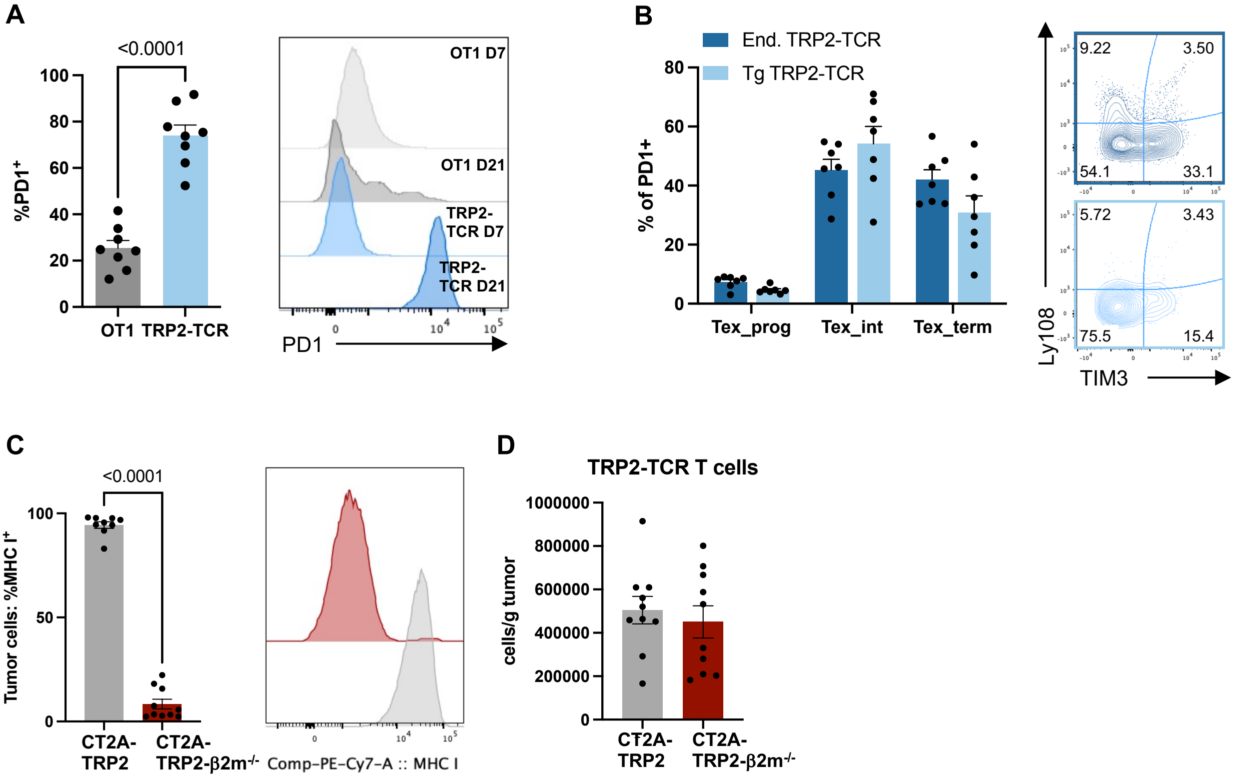


**Figure S3: Antigen-specific stimulation is required for T cell differentiation in the tumor, related to Figure 3.**
(A) PD1 expression on transferred OT1 vs. TRP2-TCR T cells on day of adoptive transfer (D7) and day of tumor harvest (D21) and representative histogram. n = 7 mice/group. (B) Frequency of Tex subsets of PD1+ endogenous or transferred TRP2-TCR T cells and representative contour plot. n = 7 mice. (C) gMFI of MHC I in CT2A-TRP2 vs. CT2A-TRP2-β2m-/- validating loss of MHC I. n = 10 mice/group. (D) Number of TRP2-TCR T cells/g of tumor in CT2A-TRP2 vs. CT2A-TRP2-β2m-/-. n = 10 mice/group. p-values are plotted on individual graphs. Statistical significance was determined by two-sided unpaired t-test (A, C, D) or 2-way RM ANOVA with post-hoc Tukey's test for multiple comparisons (B). All data are plotted as mean ± SEM.

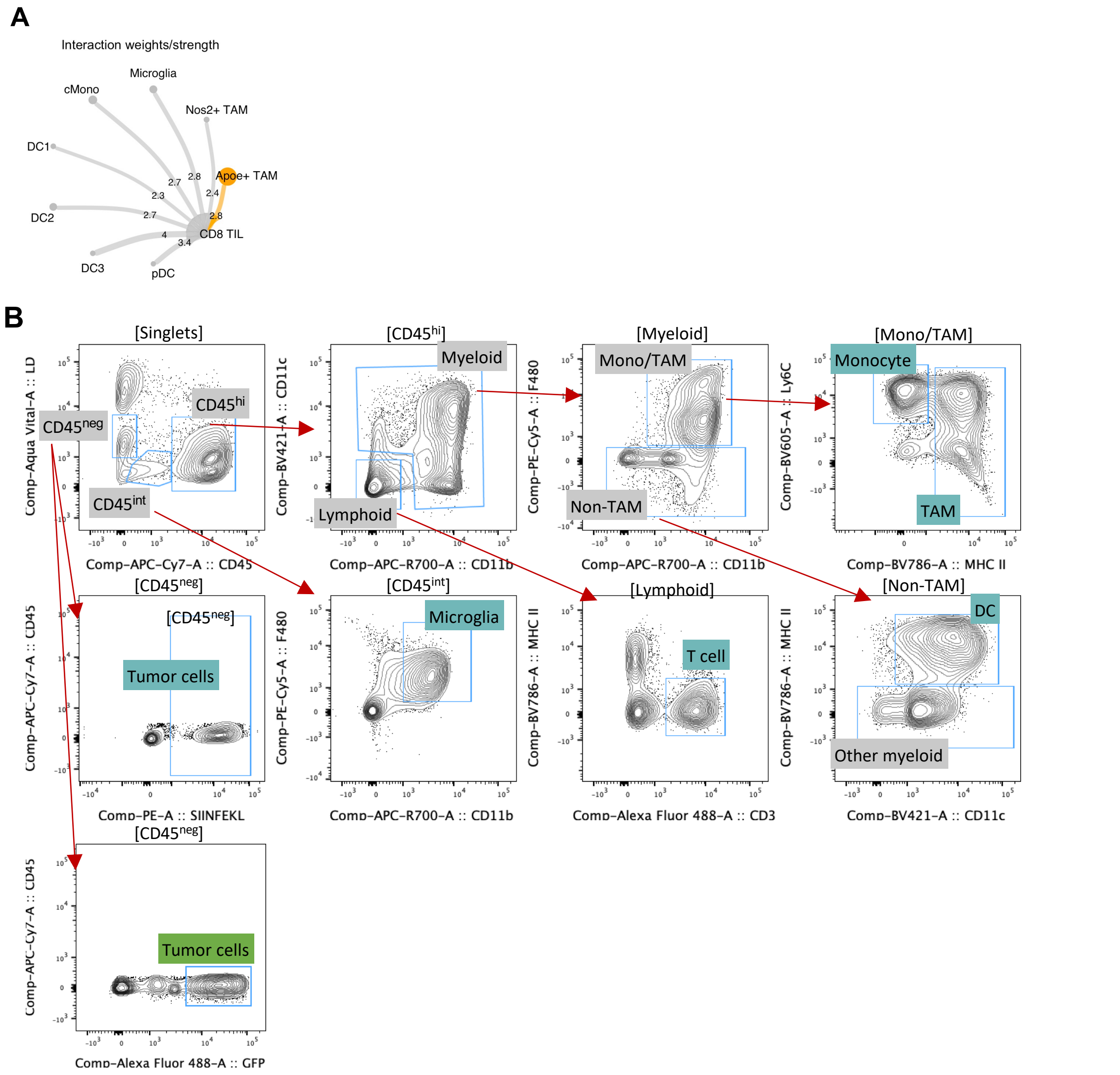


**Figure S4: Investigating APC as critical mediators of T cell exhaustion progression in the tumor, related to Figure 4.**
(A) Interaction strength across APC with CD8 TIL as the target. (B) Gating strategy for intratumoral APC.

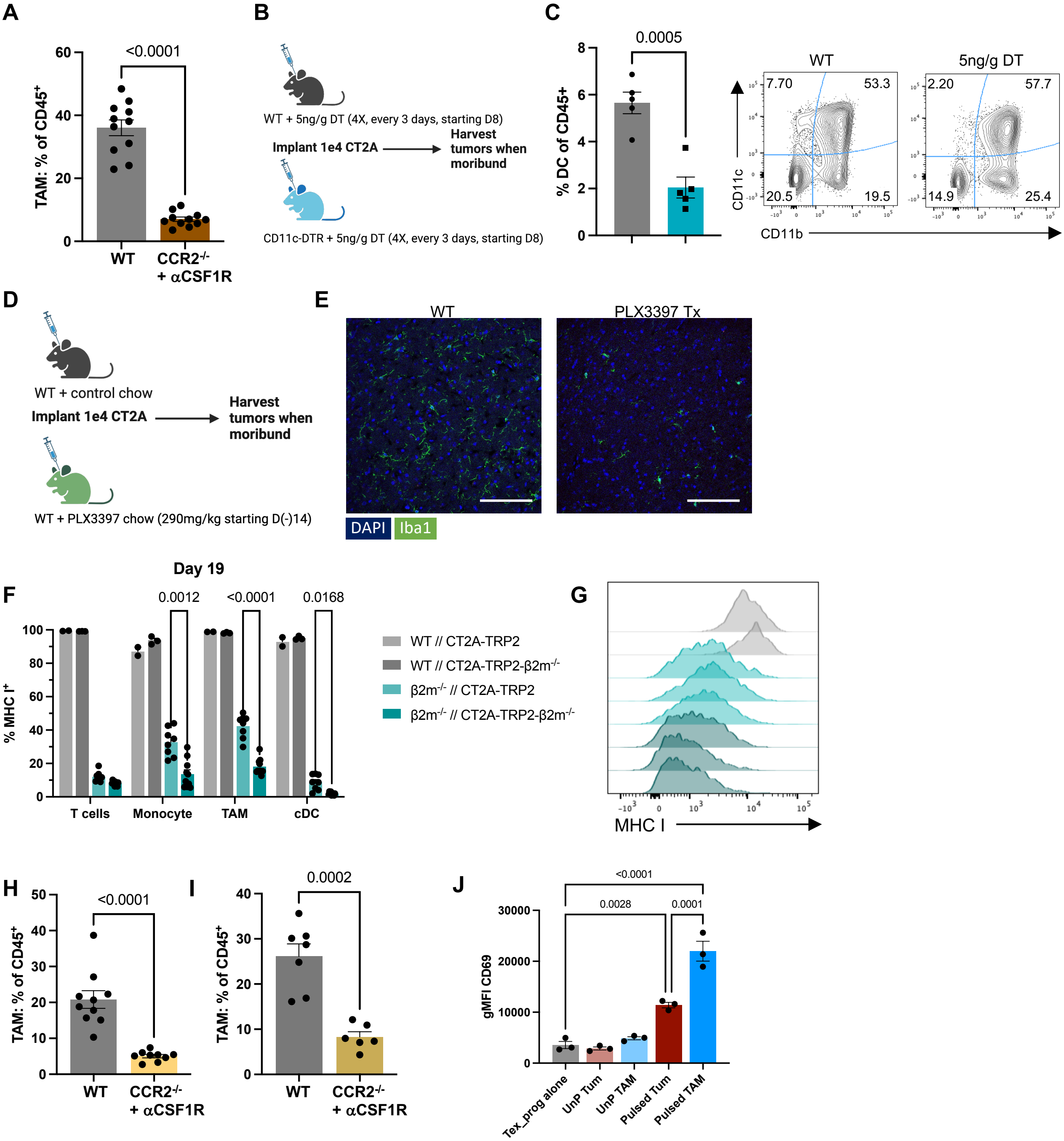


**Figure S5: Depletion of TAM, DC and microglia from glioma and melanoma tumor models, related to Figure 5.**

(A) Frequency of TAM of $CD45^+$ cells. Data pooled from 2 independent experiments. n = 11 mice/group. (B) Experimental layout. DC were depleted in CD11c-DTR mice with 5ng/g DT, every 3 days for 4 total treatments, starting D8. Created with Biorender.com. (C) Frequency of CD11c+CD11b- DC in WT vs. CD11c-DTR mice. (D) Experimental layout. Microglia were depleted in WT mice with 290mg/kg PLX3397 chow. Chow was provided starting D(-)14. Created with Biorender.com. (E) Representative immunofluorescence image of WT vs. PLX3397 treated non-tumor brain. Scale bar = 100um. (F) Chimera validation. Frequency of MHC I+ in WT → WT (WT) or $\beta2m^{-/-}$ → WT ($\beta2m^{-/-}$) chimeras across T cells, TAM and cDC. n = 8-10 mice/group. (G) Representative histogram of MHC I expression on TAM in the setting of CT2A-TRP2 (Teal) vs. CT2A-TRP2- $\beta2m^{-/-}$ (Dark Teal). (H) Frequency of TAM of $CD45^+$ cells in B16F0 IC. n = 9-10 mice/group. (I) Frequency of TAM of $CD45^+$ cells in B16F0 SQ. Data pooled from 2 independent experiments. n = 6-7 mice/group. (J) Expression of CD69 on OT1 T cells when co-cultured with TAM or tumor. n = 3 biological replicates. Data are representative of 2 independent experiments. p-values are plotted on individual graphs. Statistical significance was determined by two-sided unpaired t test (A, H, I), 1-way ANOVA with post-hoc Tukey's test or RM-2-way ANOVA with post-hoc Tukey's test for multiple comparisons (F). All data are plotted as mean ± SEM.

### Table S1: Exhaustion signature (Im et al., 2016), related to Figure 2

| |
|---|
| *Pdcd1* |
| *Tox* |
| *Il21r* |
| *Sh2d2a* |
| *Cd8a* |
| *Nkg7* |
| *Fgl2* |
| *Entpd1* |
| *Fasl* |
| *Ccl5* |
| *Apol7e* |
| *Arhgap9* |
| *Nedd9* |
| *Cd38* |
| *Pvrig* |
| *Trbc2* |
| *Chn2* |
| *2900026A02Rik* |
| *Pstpip1* |
| *Chst12* |
| *Cd3e* |
| *Cxcr6* |
| *Cd3g* |
| *Adgrg1* |
| *Prkch* |
| *Tap2* |
| *Irf1* |
| *Tapbpl* |
| *AW112010* |
| *Arl6ip1* |
| *Gng2* |
| *Adam19* |
| *Gimap7* |
| *Gzmk* |
